# Decoding of human identity by computer vision and neuronal vision

**DOI:** 10.1101/2021.10.10.463839

**Authors:** Yipeng Zhang, Zahra M. Aghajan, Matias Ison, Qiujing Lu, Hanlin Tang, Guldamla Kalender, Tonmoy Monsoor, Jie Zheng, Gabriel Kreiman, Vwani Roychowdhury, Itzhak Fried

## Abstract

Extracting meaning from a dynamic and variable flow of incoming information is a major goal of both natural and artificial intelligence. Computer vision (CV) guided by deep learning (DL) has made significant strides in recognizing a specific identity despite highly variable attributes^1,2^. This is the same challenge faced by the nervous system and partially addressed by the concept cells—neurons exhibiting selective firing in response to specific persons/places, described in the human medial temporal lobe (MTL)^3–6^. Yet, access to neurons representing a particular concept is limited due to these neurons’ sparse coding. It is conceivable, however, that the information required for such decoding is present in relatively small neuronal populations. To evaluate how well neuronal populations encode identity information in natural settings, we recorded neuronal activity from multiple brain regions of nine neurosurgical epilepsy patients implanted with depth electrodes, while the subjects watched an episode of the TV series “24”. We implemented DL models that used the time-varying population neural data as inputs and decoded the visual presence of the main characters in each frame. Before training and testing the DL models, we devised a minimally supervised CV algorithm (with comparable performance against manually-labelled data^7^) to detect and label all the important characters in each frame. This methodology allowed us to compare “computer vision” with “neuronal vision”—footprints associated with each character present in the activity of a subset of neurons—and identify the brain regions that contributed to this decoding process. We then tested the DL models during a recognition memory task following movie viewing where subjects were asked to recognize clip segments from the presented episode. DL model activations were not only modulated by the presence of the corresponding characters but also by participants’ subjective memory of whether they had seen the clip segment, and by the associative strengths of the characters in the narrative plot. The described approach can offer novel ways to probe the representation of concepts in time-evolving dynamic behavioral tasks. Further, the results suggest that the information required to robustly decode concepts is present in the population activity of only tens of neurons even in brain regions beyond MTL.

## Main

Nine neurosurgical patients (Extended Data Table 1) watched a 42-minute movie (first episode, season six of “24” TV series). Following the movie viewing, participants were tested for recognition memory by showing them multiple short clips of targets (taken from the same episode) and foils (taken from another episode of the same TV series) and were instructed to mark whether they had seen the clip (Fig. 1a; Methods). These participants were implanted with multiple depth electrodes as part of the clinical procedure for seizure monitoring and possible resection of the epileptogenic tissue. We recorded single unit activity from 385 neurons from multiple brain regions (Methods; Extended Data Table 2,3)^3,8–10^.

**Figure 1:**
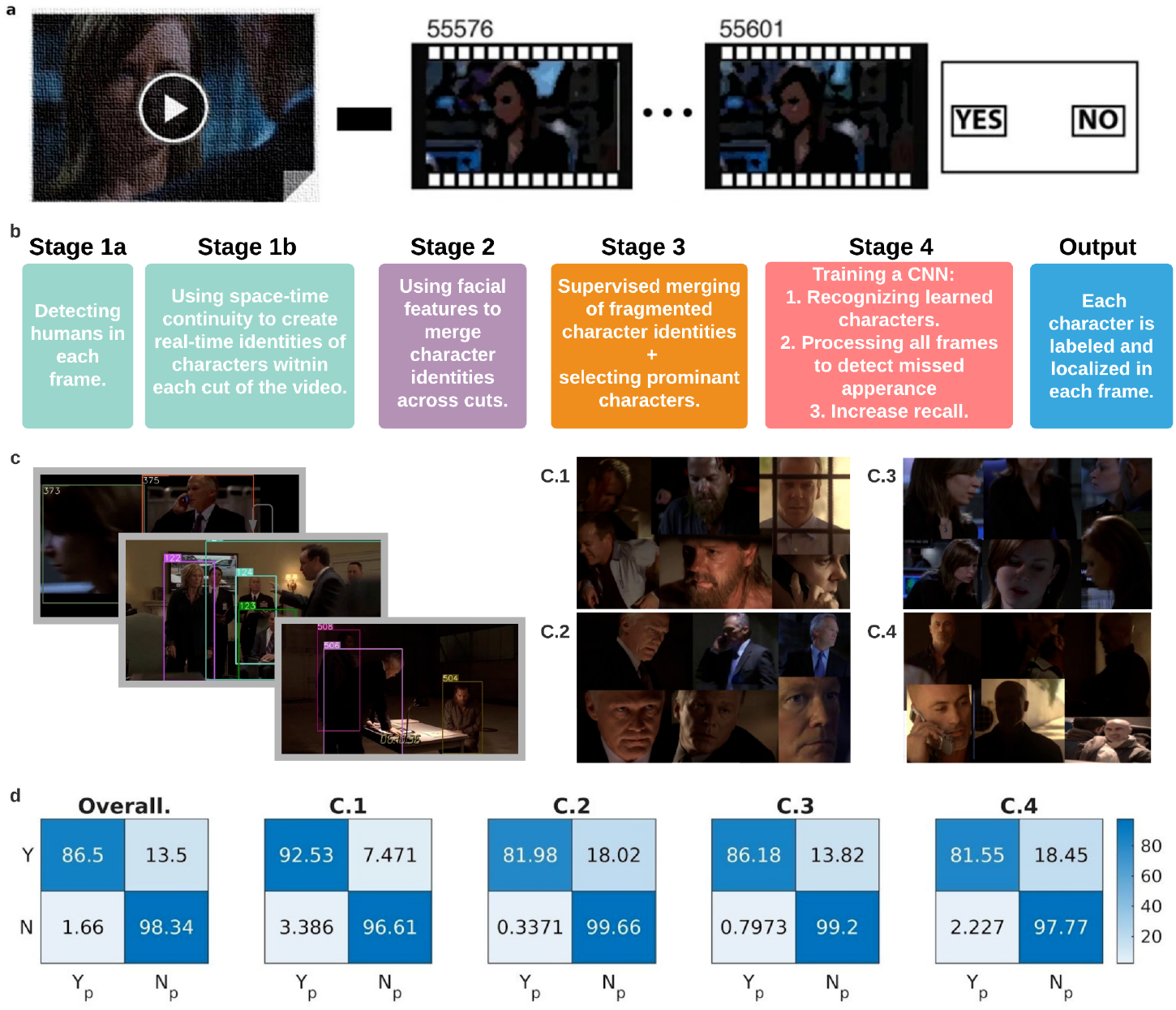
Schematic of the task and the pipeline for semi-supervised character extraction. **a.** The task consisted of viewing an episode of the 24 TV series followed by a recognition memory test. During the memory test, participants were shown short clips and were asked whether they had previously seen the clip or not. **b.** A brief overview of the algorithm used for character extraction. For a more detailed version, see Extended Data Fig. 1) **c.** Example outputs of the character extraction algorithm at different steps: Left) output of stage 1 (extracting humans in each frame); Right) output of stage 4 (sample different appearances for each of the four main characters C.1 through C.4) **d.** To test the performance of our character extraction algorithm, we used character labels that were manually created in an independent study^7^. Shown are the normalized confusion matrices of individual characters (right) as well as the overall confusion matrix (left). Yp and Np correspond to predicted yes and predicted no respectively. Note the darker colors along the diagonal showing large percentages of true positives and true negatives indicative of high performance.

We first sought to examine whether it is possible to decode the presence or absence of individual characters throughout the movie from the neuronal population responses despite the highly variable physical appearance and context. As a first step towards generating training and test sets for the decoding task, we developed a semi-supervised algorithm that labeled the presence or absence of nine important characters in each frame (Fig. 1b, c; Extended Data Fig. 1; Methods). Briefly, the associated pipeline involved: 1) extraction of humans in each frame using a pre-trained YOLO-V3 network^11^, 2) spatio-temporal tracking of detected humans to form clusters of image crops belonging to the same character, 3) grouping of these spatio-temporal image clusters into nine important character identities, based on facial features; (this is the only part that required manual intervention), and 4) training a ten-label Convolutional Neural Network (CNN) with the automatically learned examples to identify the presence or absence of each of the nine characters in each frame (the tenth label corresponded to “other”). Of the nine character identities created by our algorithm, we picked the four most prominent characters to be decoded using neural data; henceforth, to be referred to as C.1, C.2, C.3, and C.4 (This allowed us to have sufficient data points for training and testing in the later steps; see Methods; Extended Data Fig. 2; Extended Data Table 4). Prior work^7^ had segmented the same movie into a set of shots (or cuts, defined as consecutive frames between sharp transitions), and manually labeled each shot (as opposed to the individual frames in our case) with the names of the characters in it. By aggregating frame level labels over each shot, we benchmarked the performance of the automated method against the manual labels (results are shown in Fig. 1d).

Having established continuous (as opposed to a cut-level human annotation) labels for the visual presence of characters in each frame (using our semi-supervised method), we asked whether it is possible to find neural footprints for each of the four main characters in our electrophysiological data for each participant in our study. A footprint of a character is a discriminative pattern in neural data responding to the presence of the character, such that the pattern appears if and only if the character is present, and hence a decoder can be trained. Furthermore, these footprints are participant-specific, given that each participant had a unique set of recording sites. To build such a decoder, for any given frame with a character in it, we created a candidate footprint feature vector comprising the firing rates of all active neurons from all regions during a two-second interval around the frame (one second before and one second after for each target frame; see Methods). The exact number of neurons and regions for each participant can be found in Extended Data Tables 2, 3.

To implement the decoder, we first used, and optimized, a two-layer Long Short Term Memory (LSTM) network^12^ (Fig. 2a, Methods; Extended Data Table 5). The final layer outputs the probabilities of the four main characters in each frame, which during evaluation were further binarized into presence (“yes” label) or absence (“no” label) predictions (Fig. 2a). We used a 5-fold cross-validation method, with 70% of the data used for training, 10% for validation, and 20% for testing (frames were randomized and were independent in each set). It is worth noting that the true data labels were highly unbalanced and the characters were at most visually present in only around 20% of the frames, (Extended Table 4). As such, without using higher weights for the loss function corresponding to “yes” labels, one would expect that the performance would converge to a misleadingly high accuracy of 80% but would yield 0% for character detection. However, we obtained good decoding performance both in terms of accuracy and character detection, thus indicating significant latent character information in the neural data (Methods).

**Figure 2:**
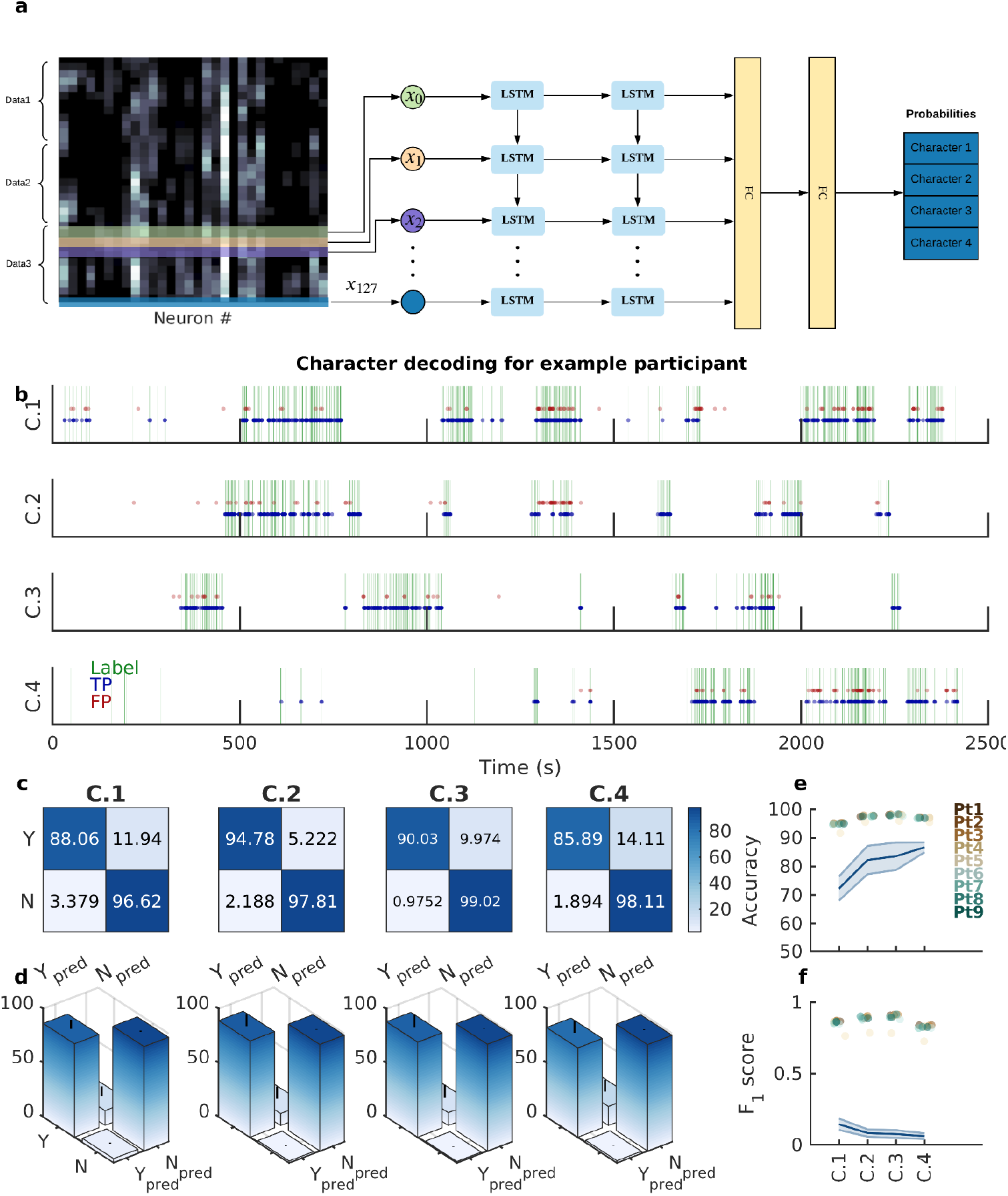
Decoding visual presence of the four main movie characters using neural recordings as inputs. **a.** The structure of the LSTM network used for classification. Here, input data was the firing sequence (colormap; brighter shades indicate higher firing) of all neurons (x-axis) from a participant within a two-second window around each frame (y-axis: time; in this case 6 seconds are shown with data1-3 example representatives of 2 seconds around each frame with 60 time-steps). Firing rate maps were sequentially passed through two LSTM layers followed by two fully connected (FC) layers to output a probability distribution over the four main characters in that frame of the movie. Note that the predictions occur on a frame by frame basis. **b.** Frame-by-Frame comparison of character labels generated by CV and neuronal vision (LSTM classifier): The labels generated by the computer vision algorithm (green vertical lines; each row is a different character) and the LSTM (blue dots: true positive, TP; red dots: false positive, FP) for each movie frame are plotted as a function of time. The significant overlap between the two labels (i.e., green lines and blue circles, large number of true positives) illustrates the goodness of the decoding algorithm. **c.** In an example participant, the normalized confusion matrices for the binary classification task for all the four characters are shown. Each row indicates the true labels (Y: character present in the frame; N: character not in the frame) and each column indicates the predicted labels by the classifier. The large numbers on the diagonals (high true positive rate (TPR) and true negative rate (TNR)) of all the four matrices shows that the LSTMs achieve high accuracy in decoding all the four characters. **d.** The distribution of the entries of the confusion matrix over all participants is shown as a bar plot (mean) with error bars (std) for all four characters. The high mean and low standard deviation for the TPR and TNR values in all the four matrices show that the LSTM achieves high accuracy in decoding all the four characters across participants. **e-f.** Accuracy (e) and F1-scores (f) for decoding each character are shown with each colored dot indicating different participants (Pt1 through Pt9). The consistently high accuracy and F1-scores across participants indicate that the LSTM generalizes well in this decoding task. The lines and shaded areas (mean±STD) indicate the performance of the chance model (obtained from shuffling labels) across all participants. Note that the chance level for accuracy is at 80% due to the unbalanced nature of the data. For instance, given that a character is present only in 20% of the frames, predicting a “N” for all frames would yield 80% correct predictions.

For each participant, the performance of the decoder was first visualized by comparing the frame level predictions for each character against the corresponding true labels (Fig. 2b), and was further quantified by a normalized confusion matrix (Fig. 2c). Performance of the LSTM decoder, as quantified by the F1-scores—a measure that is more appropriate for unbalanced datasets—was on average 7-times better than that of a distribution-based decoder (such as Naive Bayes (Methods); Extended Data Table 6). This suggests the presence of strong and unique discriminative character footprints in the neural data. The distribution of the entries of the normalized confusion matrix (Fig. 2d), the plots of the Accuracy (Fig. 2e) and F1-scores (Fig. 2f) across all nine participants, as well as the table of Recall, Precision, F1-scores, and Accuracy (Extended Data Table 7) showed consistently good results. Lastly, to further ensure that our results could not arise by chance, we performed a shuffling procedure in which the character labels were randomized with respect to the neural data and the participant-specific models were retrained and re-evaluated. Here, too, the performance of the true model was far above the performance of the chance model (Figs. 2e,f; shaded region). Next, we addressed the effect of faulty labels in the output of the semi-supervised computer vision algorithm on the performance of the neural decoder (“neuronal vision”). We used the cut-level human annotations as the ground truth. Although the neural decoder was trained and tested using the computer vision (CV) labels, which albeit close to the ground truth (manually-labelled data) contained a small number of faulty labels (Fig. 1c), we found that in the case of recall, neural vision outperformed the computer vision results (p=5.05×10^−3^, Signrank test).

Since Neural Network (NN) architectures, such as the LSTM, have high representational capabilities, several different converged models (corresponding to different minima for the same training data) could give comparable end-to-end performance but could produce dramatically different results when the models are used to determine functional properties of the underlying physical systems. In our case, for example, we intended to use the learned models to determine the regions and subregions carrying the most relevant information in the decoding of different characters. Therefore, we used an entirely different NN architecture, namely a convolutional neural network (CNN) model, where the time series training data around each frame was converted into an image (Methods; Extended Data Table 8). The exact same tasks were replicated for both LSTM and CNN networks. Indeed, the CNN model reached comparable high performance to that of the LSTM model in decoding characters (Extended Data Fig. 3; Extended Data Table 6) and yielded similar results to the LSTM pipeline in the subsequent analyses detailed below. The consistency of results between the two NN models (LSTM and CNN) is critical, especially when assessing the importance of different brain regions in the decoding process, since it ensures that the results are not merely an artifact of model optimizations.

Thus far, we used the activity of all of the recorded units within each participant as the input to the NN models. Next, we used a knockout analysis to determine the brain regions that were more critical than others in the decoding process. This knockout analysis is analogous to the analysis tool named Occlusion Sensitivity^13^ for inspecting NN image classifiers. Specifically, we evaluated the performance of our model (that was trained on units from all regions; base results) on data in which the activity of units from individual regions was eliminated one at a time (region knockout results). We used the change in the Kullback–Leibler divergence (KLD) loss due to the knockout, normalized by the number of neurons, as a proxy for how worse (or better) the model performs without units recorded from a specific region (Methods). We found that knockouts of different regions led to different changes in KLD loss, and we determined important regions to be those which, when knocked out, led to higher normalized KLD loss (Fig. 3a). Of the eleven regions, knocking out five of them resulted in the most notable (normalized) losses in decoding performance (occipital, entorhinal cortex, parahippocampal, anterior cingulate, and superior temporal)(Fig. 3b). For each participant, we additionally verified this finding by re-training two independent NNs on neural data from important and less important regions and by comparing their performance. The two separate models, one trained only using the units from regions that were deemed important, and another one trained only using the remainder of the units (Methods), showed a significant difference in their decoding performance (p=0.01, Wilcoxon ranksum test comparing the F1-scores of the two re-trained models).

**Figure 3:**
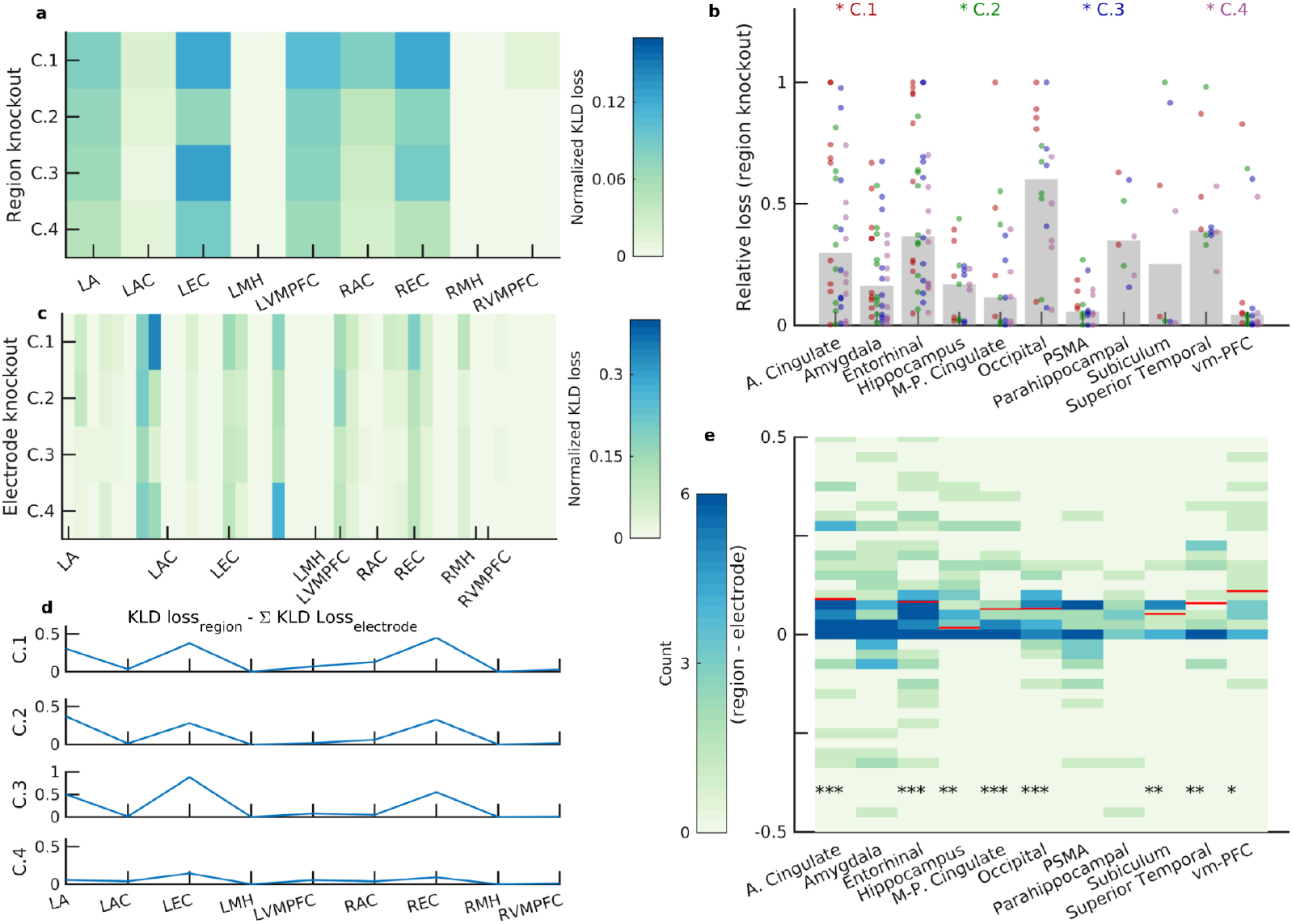
Identification of important regions in decoding characters: “The Whole Is Greater Than the Sum of its Parts.” **a.** The change of KLD loss for each character (row) after knocking out a given region (column) for one participant is shown (Region Knockout). The value is normalized by the number of neurons in that region and demonstrates how the model performance deteriorated when excluding the units recorded from that region. Important regions are those with higher KLD loss values. L and R correspond to the left and right hemisphere, respectively. A: Amygdala, AC: Anterior Cingulate, EC: Entorhinal Cortex, MH: Middle Hippocampus, VMPFC: Ventro--medial Prefrontal Cortex. **b.** The changes in KLD loss after knocking out regions are shown across participants. Different colored dots correspond to the changes in KLD loss for different characters. Bars indicate the median value of the change in KLD loss after region knockout. The following regions resulted in the most notable losses in decoding performance: anterior cingulate (36.11, [20.82, 53.78]%), entorhinal cortex (42.50, [27.04, 59.11]%), occipital (65.00, [40.78, 84.61]%), parahippocampal (37.50, [8.52, 75.51]%), and superior temporal (33.33, [9.92, 65.11]%). Reported are the percentage of losses above 0.5 (as well as the binomial fit confidence intervals).(N_participants_=9). **c.** The change in KLD loss for each character (row) after knocking out a given electrode (column) at a time is shown (Electrode Knockout) for an example participant (same as in a). Similar to the region knockout results in (a), the loss value is normalized by the number of units recorded on each electrode. **d.** The sum of the changes in KLD loss following electrode knockout (all electrodes within a region) was subtracted from the change in KLD loss following region knockout. Shown are these values for the four different characters (rows) from an example participant (same as in a and c). Positive values indicate that knocking out a whole region deteriorates the model performance to a greater extent. **e.** When considering all regions from all participants, in most regions, the region knockout loss was greater than the sum of electrode knockout loss (each column, and its associated colormap, is the distribution of this measure and the red horizontal line indicates the median of the distribution for those that were significantly different from zero) as quantified by Wilcoxon signed-rank tests (*: p<0.05; **: p<0.01; ***: p<0.001).

We asked whether the co-activation pattern among the neurons within each region was contributing to the decoding performance. We defined the incremental information content of a neuron as the increase of the KLD loss on removing the neuron’s activity while preserving the rest of the system. We observed that the information content of a set of neurons, that is, the increase in the KLD loss by removing all the neurons in the set, is larger than the one obtained by summing up the incremental information contents of the individual neurons. This was shown by applying the same knockout analysis, where we knocked out the activity of all the neurons recorded on individual electrodes (microwires; 8 per region; electrode knockout results) one at a time and evaluated the KLD loss (Fig. 3c). Our analysis showed that the resulting increase in the KLD loss from region knockout (i.e., when all the units in a given region were knocked out together) was greater compared to when the increases in KLD losses from electrode knockout within a region were added together (Fig. 3d) in most regions (Fig. 3e; P<0.05 for eight out of eleven regions, Wilcoxon signed-rank test). These results were replicated using the CNN network as well (Extended Data Fig. 4). This finding of “the whole is greater than the sum of its parts” may indicate that the neurons’ dynamics and inter-relations across a region may also contribute to the decoding performance.

After quantifying the model performance during movie viewing, we examined how the model fared during the memory test following the movie (Fig. 1a; Methods). It should be borne in mind that because of the nature of the recognition task—where participants have to decide whether they have seen the clip or not—participants would most likely remember parts of the movie plot beyond those displayed during the clips. Thus, any well-trained decoder may predict the presence of characters not necessarily visually present in the clip itself. As such, when any such decoder is evaluated by whether it predicts the characters in the clips, it might lead to more false positives (FPs) compared to the movie viewing time, therefore lowering the accuracy. Indeed, as reported in the following, we observed that our decoder led to an increased number of FPs and, as expected, the accuracy of our model was lower during the memory task compared with the movie (~67% compared to ~95%). We discovered, however, that the accuracy of the model was positively correlated with both the percentage of the time the character was present in the clip, as well as, the size of the character in the frames (Fig. 4a), which might be expected given that both are measures of how prominent the character is.

**Figure 4:**
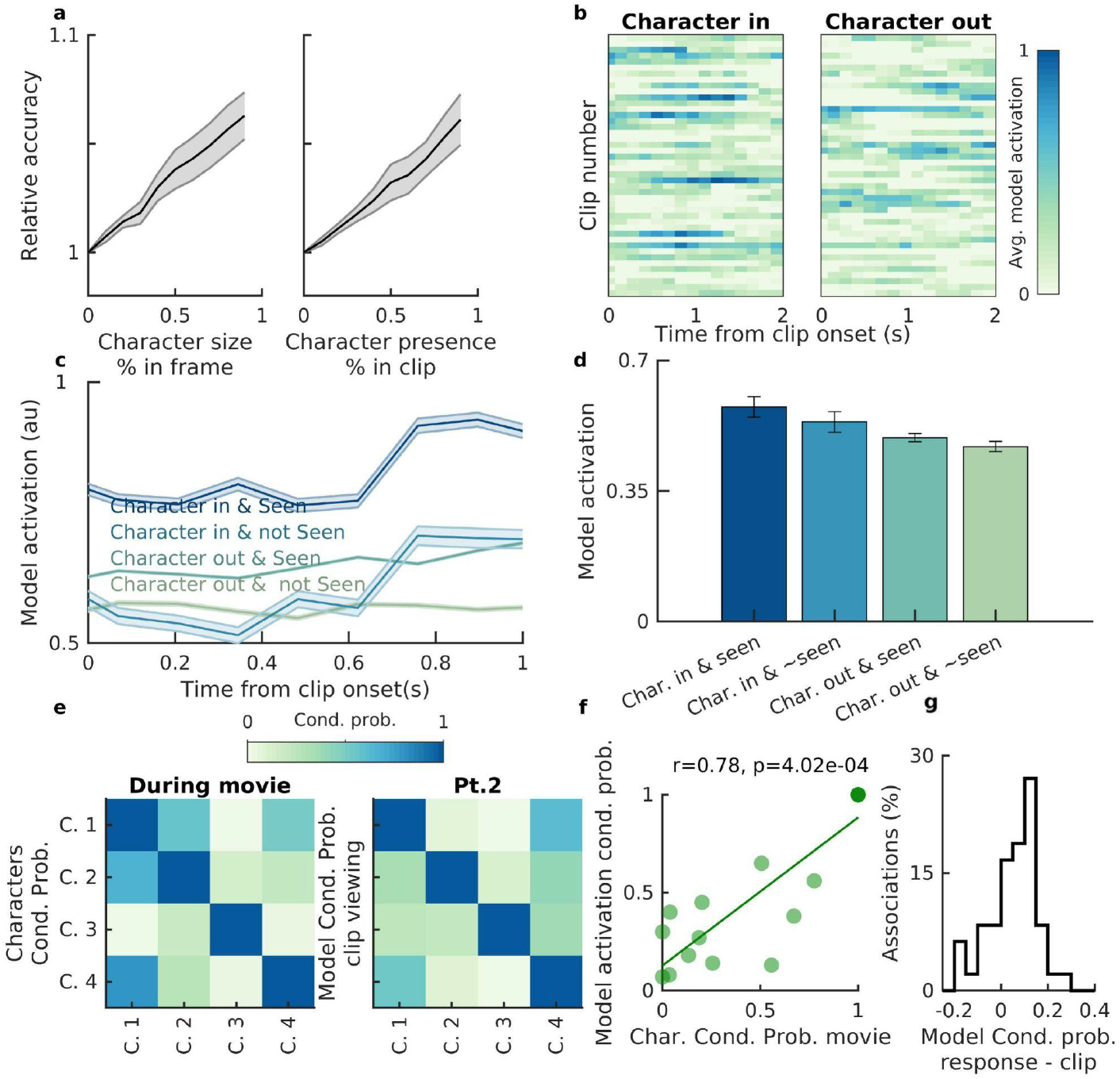
Properties of the NN model trained for decoding characters in the movie during the memory test. **a.** The accuracy of the NN model (trained during movie viewing) in decoding the visual presence of the characters in the clips during the memory test improved as a function of both the “size” of the character in the clips (defined as the percentage of the pixels that the character occupied in a given frame; reference: half frame) as well as the prevalence of the character in the clip (defined as the percentage of the clip time during which the character was present). Shown are the mean±SEM of the model accuracy at different thresholds across participants. **b.** Model activation as a function of time (averaged across five folds) was separated for the clips during which the character was present (character in; left) and all other clips without the character (character out; right; a random subset of the clips are shown in the panel). Note that during the clips that contained the character the model activation appeared higher. The example shown is from participant 5, activation for character 1. **c.** Model activation as a function of time (with respect to the clip onset), was divided into a 2×2 matrix: clips with/without the characters x participants’s subjective memory of the clip (yes-seen/no-have not seen). Shown are the mean±SEM of the model activation for each of the four groups of clips. d. To test whether both the presence of characters and participants’ subjective memory influenced the model activation at the population level (for all participants and clips), we used a GLM method. We modelled the overall activation (during a two-second interval after clip onset) as a function of 1) whether the character was in the clip or not (estimated coeff.=0.29, p=9.29×10−4); and 2) whether the participant marked the clip as previously seen or not (estimated coeff.=0.15, p=0.03). Both were significant factors in the model activation and the height of the bars and error bars correspond to the mean overall activation and the standard error across clips (from all participants) respectively. **e.** Left) Character associations as quantified by the conditional probabilities of their cooccurrences during the movie. For example, row 1, column 4 represents P(character 4 | character 1). Right) Conditional probabilities of model activations for the characters. The structure follows the same pattern as in (left). **f.** Conditional probabilities of model activations during clip viewing (e-Right) were significantly correlated with character associations (e-Left) in an example participant (Spearman correlation r=0.67, p=4.73×10^−3^). Please note that the values from the diagonals are excluded in this analysis). **g.** Conditional probabilities of the model activation for characters were higher during response time compared to clip viewing (p=0.016; sign-rank test).

Our NN models output a confidence level for each character in the range [0,1] at any given time; this confidence level is henceforth referred to as model activation. So far, while reporting metrics such as accuracy and F1-score, we followed the standard technique of binarizing this model activation: 1 if it is greater than 0.5 and 0 otherwise. However, model activation can provide more granular information and was, therefore, used to analyze the decoder performance during the memory task. We evaluated the NN model activations for each character as a function of time and, specifically, with respect to the clip onset time (Fig. 4b). We noted that the model activations, in addition to the visual presence or absence of the characters, were also related to participants’ subjective memory. Model activation during clip viewing was highest when the characters were in the clip and the participants later marked the clip as “seen before” during the response time. Conversely, model activation was lowest when the character was not in the clip and the participant marked the clip as “not seen before” (Fig. 4c). At the population level, both the presence of the character as well as the participants’ response to whether they had seen the clip were significant contributing factors in model activation as revealed by Generalized Linear Models (GLM) (Fig. 4d; Character in: estimated coeff.=0.39, p=8.02×10^−6^; Clip marked as seen: estimated coeff.=0.24, p=0.001; GLM estimates). The effect of participants’ subjective memory of the clips on the model activation during memory was surprising given that the NN model was only trained to distinguish the visual presence or absence of the characters. When we performed the knockout analysis (described in the previous sections) and evaluated the model activations after knocking out the activity of all MTL neurons, participants’s subjective memory was no longer a significant predictor of the activations, i.e. only the character presence remained a significant factor (GLM estimated coeff. = 0.50, p=9.50×10^−9^), which is consistent with the role of the MTL in the formation of memories^14–16^.

It has been shown previously that the formation of new associations are reflected in the firing pattern of single neurons^9^. Accordingly, we hypothesized that the emergence of NN model coactivations would reflect character associations. To test this, we quantified two measures in parallel: 1) character associations: we utilized a segmentation of the movie into a set of scenes (provided by an independent study^7^) where each scene represents a meaningful subplot that is expected to create associations in viewers. We computed the conditional probabilities of the characters over the scenes (e.g., the probability of character 1 appearing given character 2 appears in a given scene (Methods), henceforth referred to as character associations; Fig. 4e); 2) NN model coactivations: We noted that model predictions for characters during clip viewing exhibited overlapping activations amongst the characters. For example, when the model activation was high for character 1, there was a higher activation for character 4 as well (even in the absence of character 4, which led to lowering the general performance of the model in decoding characters). Thus, we computed the conditional probabilities for the characters predicted by our model in the clips (e.g., the probability of model activation being above 0.5 for character 1 given the model activation is above 0.5 for character 2)(Methods). We found that the conditional probabilities of the model activation for characters during clip viewing were positively correlated with character associations in the movie (example: Fig. 4f; for all participants: p<0.05, Spearman correlation).

Lastly, we inspected the model activation during the response time (following the end of the clip and prior to the participants making a choice) when no visual information was provided to the participants, and noted that conditional probabilities of model activation for the different characters were significantly higher compared to those during clip viewing (p=0.016; sign-rank test; Fig. 4g), which might suggest higher-order associations among the characters rather than first-order associations as observed during clip viewing, may be invoked during the response time.

## Discussion

We report on an integrated methodology that allowed us to perform neuronal decoding of individual characters, each with highly variable sensory and contextual depiction viewed throughout a 42-minute audiovisual sequence, and to compare it to the ability of computer vision with minimal human supervision to recognize these characters’ identities. On the one hand, we proposed a semi-supervised algorithm to extract the characters in each movie frame to achieve continuous labelling of data for further analysis. On the other hand, we implemented a DL-based framework to decode the visual presence of characters using neuronal data. Our NN model further allowed us to extract and follow the characters’ footprints in time and investigate how they interact during memory tests.

Our semi-automated method served two purposes: first, its performance was comparable to that of human labeling, and thus it could be used as a general tool for future research in this field without the need for time-consuming and expensive use of Mechanical Turk resources; second, it showcased what state-of-the-art computer vision technology stack can achieve (along with minimal human intervention) if it were to watch the movie itself instead of a human. In particular, whether it could accurately create visual models of different character identities, and then recognize them in every frame. The full power of CV tools provided an interesting contrast to the information contained in the firing patterns recorded from only a few tens of neurons.

Our ability to decode the visual presence of the characters using the neural data, even in the absence of direct access to concept neurons explicitly representing these characters^3,6^, suggests that there is indeed sufficient information in relatively small populations of neurons, even in regions beyond the MTL, to detect the character footprints. Given that we obtained similar results—in both decoding of the characters as well as identifying the important regions contributing to the decoding process—using two different neural networks, it is unlikely that our results are merely artifacts of the trained networks. Of note, the sensory and association regions—such as the occipital and superior temporal—contributed most to the decoding process^14–16^. In addition to highlighting the brain regions that may be recruited in the representations of the characters, this approach suggested that the correlations amongst the neurons within a region may be conducive to the decoding process. This observation perhaps resembles the information theoretic phenomenon of synergy^17^ where the aggregation of knowledge from multiple sources results in more information than the sum of the sources.

Despite having trained the NN models to only decode characters during movie viewing, the model performed well during the memory test. Although, admittedly, the model performance during the memory test was slightly lower compared to movie viewing. This lower NN model performance could be potentially explained by the changes in neural representations due to the change in context (viewing versus remembering) or the formation of composite representations due to forming associations with time. The model activation during the memory test seemed to corroborate previous hypotheses on memory reinstatement given that the participants’ subjective memory was, in addition to the presence of characters, a significant predictor of the NN model activations. This may suggest that the patterns of neural activity during movie viewing might be reinstated^18–22^ differently in accordance with participants’ memory of whether they had previously seen the clip or not, which consequently affects our NN model predictions—an effect that was eliminated when the activity of MTL neurons were discarded from the models.

Moreover, during the memory test, the model predictions may be capturing what has been conjectured regarding the change in the neuronal representations as a result of forming associations between items^9,23^. How the model coactivation for the characters during the memory test reflected the relationship between the characters, and their associations, in the movie itself may be reminiscent of associative coding schemes. Although our knockout analysis to examine whether this effect was specific to the medial temporal lobe regions or not did not lead to convincing results, examining the role of MTL vs. non-MTL neurons in the representation of associations warrants future investigations with large-scale data.

Taken together, our proposed platform offers an approach to interrogate the neural signatures involved during a temporally continuous experience that is aimed at mimicking real-life episodes and going beyond the stimulus-response experiments. Furthermore, it allows inquiries beyond the possibility of character decoding and shed light on potential mechanistic insights into memory processes.

## Acknowledgments

This work was supported by the NIH NINDS (NS033221 and NS084017 to I.F.) and NSF Center for Brains, Minds and Machines and McKnight Foundation (G.K.). We also thank the participants for taking part in our study.

## Materials and Methods

### Participants

Nine epilepsy patients (6 Females, 20-50 years old; Extended Data Table 1) who were implanted with intracranial depth electrodes for epilepsy treatment (seizure monitoring) participated in our study during their hospital stay. Prior to the surgery, an informed consent was obtained in accordance with the Institutional Review Board at UCLA.

### Behavioral Tasks

Nine participants watched an episode of the TV series 24 (season 6, episode 1) and were tested for recognition memory following the movie viewing.

The memory recognition task consisted of participants being presented with short clips. Half of the clips, chosen randomly, were from the episode they had just watched (target clips), and the other half were chosen from the second episode of season 6 (lure clips). Because the episodes occur in two subsequent hours of the day, the characters look similar between the two episodes, and hence the clips from the second episode seemed appropriate as lure clips (for further justification, see *ref. ^7^*). After the end of each clip, participants were asked to make a choice of whether they had seen the clip before or not. Thus, the memory phases consisted of interleaved “clip viewing” and “response time” sections and the number of presented clips varied from participant to participant (range: 100 - 300 clips).

### Data Acquisition

Electrophysiological data were recorded from the microelectrodes located on the depth electrodes (placement was determined by clinical criteria). Wideband Local Field Potential was recorded using a 128-channel Neuroport recording system (Blackrock Microsystems, Utah, USA) sampled at 30 kHz.

### Neural data preprocessing

Previously used methods were used for spike detection and sorting^3,8–10^. A bandpass filter in the range 300-3000Hz was applied to the broadband data to detect spikes. Spikes were first automatically sorted into clusters using the Wave_clus toolbox and the clusters were further manually inspected for: 1) spike waveforms; 2) refractory spikes; and 3) the interspike interval for each cluster. The units with a mean rate below 0.05Hz were discarded for further analysis.

Additionally, we constructed a spike train by temporarily binning the spikes into 100ms bins. We further visually inspected the data, a matrix of the size NxT, where N was the number of units and T was the number of time bins. Time bins with synchronous activity across multiple regions were deemed as artifacts and were excluded from further analysis.

After data preprocessing and visual inspection, the cleaned spikes in selected regions were binned into 20ms bins. Then the spike counts were interpolated linearly into 15Hz signals.

### Electrode Localization

Electrode localization was done by co-registration of a high-resolution post-operative CT image to a pre-operative whole brain and high-resolution MRI for each participant using previous methods^10,24^ (Extended Data Table 2).

### Semi-supervised character identification

We developed a semi-supervised framework for identifying the characters in the movie at the frame level. The original movie was sampled at 30 frames per second and to make the framework computationally efficient, we downsampled the frames by a factor of 4, i.e. every fourth frame was retained. After downsampling, the number of frames in the 42-minute movie was reduced to 18900. The semi-supervised framework consists of three main stages, which we described below:

1. *Stage 1:*

a. We utilized a common practice in filming, where a particular visual setting is used to describe a part of the underlying plot. This visual setting is characterized by a distribution of colors and other features in the frames which are distinct from those of the preceding and succeeding settings, allowing one to automatically split the video into segments, *which we called* ***cuts***. We used PySceneDetect^29^ to generate such cuts. Each such cut, represented by a sequence of frames (clip of the video), will be our unit of analysis. The primary advantage in extracting such cuts is that during each cut, a group of mostly unchanged characters play their roles, thus, allowing us to create real-time identities of these characters via local tracking and similarity metrics, as described next.
b. We processed each cut separately and created real-time identities of distinct characters as follows. (i) We first used a pre-trained YOLO-v3^11^ to detect humans and drawed the bounding boxes of them in each frame. If the bounding boxes were too small or the aspect ratio was highly skewed, we filtered them out (thresholds are specified in the code accompanying the manuscript). (ii) Then we used SORT^25^ to group all bounding boxes belonging to the same character in one group, solely based on the principle of spatio-temporal continuity: each character will follow a unique trajectory across consecutive frames; no discriminating image features are used. The framework uses two algorithms, (a) A Kalman filtering tracking to follow the bounding boxes across frames; however, when characters cross each other (in the 2D frame space) or when a new character appear, the trajectories cannot be disambiguated by tracking alone; (b) A Hungarian matching algorithm to resolve such ambiguities by assigning the correct labels. For example, if the 2D projections (as captured by frames) of two 3D trajectories intersect and then separate into two, the Hungarian matching algorithm can label one outgoing trajectory as the continuation of the correct incoming trajectory by matching the bounding box dimensions before and after the intersection. For each such distinct trajectory, we collected and cropped (from the original frame images) all the bounding boxes belonging to it. These image patches will be referred to as *crop clusters*. Thus each cut will generate multiple crop clusters, corresponding to the number of significant characters in that cut. There are two kinds of errors that might be generated in this stage: 1. Multiple crop clusters within the same cut might correspond to the same character. 2. Several characters might show within one crop cluster. We manually checked the result in this stage, and the mistakes were found to be negligible. At the end of this stage, we got 1705 crop clusters and each crop cluster is pure in the sense that the image patches belong to a single ground-truth character.
2. *Stage 2:*

a. We merged the crop clusters belonging to the same character across all the cuts purely based on facial features. This grouping of the crop clusters was done iteratively, using k-means clustering and KNN. The feature vector of crop cluster Ci was extracted in the following manner: i) Each image in cluster Ci was fed into a pre-trained FaceNet^26^ to extract a 128-dimensional feature vector from the last layer of the network. Note that if no face is detected in a given image patch, we do not use this image’s patch in this stage. Moreover, if any crop cluster loses 80% of its image patches due to lack of facial features, we removed this crop cluster. The attribution of such image patches dropped in this phase to characters is made at Stage 3 using the trained ResNet. ii) Averaging the feature vectors of all of the remaining image patches in this crop cluster Ci produced the final feature vector F(C_i_).
b. The second step consisted of first clustering crop centers using k-means, yielding k super-clusters. Each such k-means super-cluster was evaluated using a distortion metric. A good super-cluster with low distortion was retained defining a character; note that multiple super-clusters at this stage can represent the same character. For example, if feature vectors F(C_i_)’s of crop clusters C_1_, C_2_… C_j_ comprise a good super-cluster (i.e. with distortion metric below a threshold) after k-means, then image crops belonging to all these crop clusters were merged together into a single group (which we referred to as a ***supernode***) representing the visual snapshots of one unique character in the video. The bad super-clusters after k-means were disintegrated into their constituent crop clusters: each such crop cluster that belonged to the bad super-clusters will be evaluated to see if they can be merged with a good super-cluster and hence it will be referred as a candidate crop cluster (CCC). Since the clustering results depend on the choice of k (the number of clusterings) in the k-means algorithm, we picked a relatively large k to balance the purity and completeness of supernodes (good super-clusters). This choice of a large value of k ensured that bad super-clusters always existed, allowing the supernodes to be pure.
c. Next, realizing that the bad super-cluster may have crop clusters that would otherwise be a good match for the supernodes, k-nearest neighbors algorithm (KNN) was used to determine such crop clusters and assign them to match supernodes. This increases the sizes of the supernodes, improving coverage while retaining purity. We defined image distance between two image patches as the euclidean distance of their feature vectors extracted from FaceNet. Additionally, we define cluster distance from crop cluster A to crop cluster B as the median of the collection of K nearest image distances from each member in A to those in B. Then we define the distance from a candidate crop cluster (CCC) to a supernode as the minimal distance from the candidate to any crop cluster in the supernode. Then the assignment of the candidate crop cluster (CCC) to a particular supernode is determined by the smallest distance among all supernodes. If the distance to the winning supernode is below a threshold, then this candidate crop cluster is assigned the same label as the winning supernode. To summarize, crop clusters with the same pure character were first grouped into supernodes by k-means and then image level feature comparisons were used so that the supernodes still remain pure while absorbing candidate crop clusters.
d. At the end of this KNN operation, we had supernodes (with absorbed candidate crop clusters) and isolated crop clusters that were not merged into any of the supernodes. We defined this one pass of k-means clustering and KNN reassignment as one iteration. After each iteration, we computed the increase in the size of each supernode. If the increase in the size of any supernode is above a threshold, implying there is potential to still absorb more crop clusters, then we move to the next iteration where each supernode is treated as a single crop cluster. Otherwise, we moved to the third step in this stage. In our experiment, we achieved satisfactory results after two iterations. After the first k-means, we had 21 supernodes and over 600 bad candidate crop clusters. Then around 100-200 bad candidate crop clusters were merged into supernodes in each iteration. After two iterations we generated 32 supernodes with consistent character appearance within each supernode and leftover 300 isolated candidate crop clusters.
e. The clustering obtained after the end of the above processing might not still be perfect because the same character can be in different clusters. We fine-tuned the clusters to further increase the completeness of clustering in the following manner: i) Constructed a fully connected weighted undirected graph with the clusters as the nodes and the euclidean distance between the cluster centroids as the edge weights ii) Pruned the network using frame distribution and edge weights iii) Extracted the connected components of the network as the fine-tuned clusters. Finally, we obtained 305 fine-tuned clusters having one character in each of them and used human supervision to aggregate and select 9 major characters.
3. *Stage 3:* We obtained 305 fine-tuned clusters having one character in each of them and used human supervision to aggregate and select 9 major characters which comprised 200 clusters. We manually inspected each cluster and assigned them character labels (e.g., clusters of the same character), see Extended Data Fig. 5.
4. *Stage 4:* Until now, we were able to recognize characters in most generated crop clusters from original cuts in stage 1. However, only frames with high quality facial features are labeled in this processing. We intend to improve coverage using data augmentation from spatial-time correlation. To be more specific, based on the time and space continuity in the cuts, the label is augmented by assigning the same label to the back view or blurred images as the clear and frontal view faces in the same cut. At the same time, after stage 2, we dropped the whole cuts with only back view or blurred characters, which are not further processed in the grouping and human-labeling. To recall those character occurrences in the original video, we trained ResNet^27^ using augmented labels for those human selected nine characters. We expected the trained ResNet to successfully classify the images according to its generalization ability. The trained ResNet was used to detect characters in all the crops from stage 1. As a result, the number of recalls of each character increased around 20% compared to that from stage 3.

The above-mentioned procedures were initially implemented on the first episode and the reported numbers are associated with episode 1. Subsequently, we applied the same method on the second episode to detect the characters in each frame since during the memory test the lure clips were taken from episode 2.

### Comparison of our semi-supervised method against human labeling

To validate the performance of our pipeline, we compared our results against manual character labeling of the movie that was done in an independent study^7^. The independent study divided the whole movie into 1001 cuts (segments between two sharp transitions) and marked which characters were present during each cut (even when present only for a subsample of the cut). Since the manual labeling was done on these cuts, as opposed to individual frames, we upsampled our character detection results to accommodate the cut base. We introduced a filter to further denoise our predictions. In one cut defined by the independent study, we viewed any character as appearing in that cut if the said character was identified for at least 12 frames in the cut. If the cut was longer than 1.3 seconds, for a character to be present in that cut, the detection of that character had to be more than 40% of the frames. Otherwise, the character had to be detected in at least 12 frames.

We then computed the confusion matrix for our model predictions against the human annotations of the four dominant characters, and the results are reported in Fig. 1d.

### Deep Neural Network (NN) implementation for neural data processing

NN implementations were done using Python and the PyTorch framework. This procedure consisted of multiple sections that are detailed below:

1. *Training data sample generation (inputs):* The training dataset of the NN model was generated by sampling the neural signal (firing rate sampled at 30Hz) around the timestamp of each frame. Since the total number of frames used in character identification is 18900, we created 18900 corresponding windowed neural signals as training data with character occurrence descriptors as labels. Concretely, for each frame, we retrieved neural data one second before and after as one training data point. Therefore one training data is a 2d matrix with shape 60 * number of neurons (in each participant). It is worth noting, however, that this duration interval of two seconds around each frame was determined by minimizing the length while meeting good decoding performance.
2. *Training data sample generation (outputs):* The corresponding label for each training data sample was constructed by three categories: Yes, No, and Do-Not-Know (DNK) for each character. Unlike traditional binary classification tasks, we introduced a third category for each character to handle the ambiguity of the visual signal. Yes was defined as when the character was present in a given frame, and No was defined as when the character was not shown in a given frame with certainty. DNK is a mechanism to particularize the training set. If an individual character is located in the predefined ambiguous time intervals (for example, when two characters are having a dialogue in a sequence of frames but some intermediate frames include only one character at a time), we allocate a DNK label to the temporarily absent character for those frames. However, this data sample could still be used by other characters in the decoding task. The percentages of Yes and No labels were around 10% and 88% of the total data samples, respectively (exact values are reported in Extended Data Table 4), showing that characters’ appearances are not evenly distributed. Moreover, we took measures to accommodate the skewed nature of our training data when quantifying the performance of our models (see 7. *Further quantifications of the model performance*). The percentage of DNK labels were around 2% of the total training samples, which indicates that the dropping of these data samples would not affect the total information provided to the model. The class DNK is discarded in all of the visualizations and performance analyses.
3. *LSTM:* A suitable type of network for our purposes was the Long Short Term Memory (LSTM)^12^, which has been already employed in neural signal processing^28^ due to its ability to extract patterns from sequential data. The architecture of the LSTM network is shown in (Fig. 2a, Extended Data Table 5). The raw windowed neural signal was fed into the LSTM directly. We conducted several preliminary evaluations to optimize the best network architecture by varying the number of hidden layers and their size (i.e., the number of LSTM units for each layer). The best network was found to have two hidden layers with 128 LSTM units each. The last layer of the hidden state of the LSTM was fed into two sequential Fully-Connected layers with LeakyReLU activation and Batch Normalization in between. The output of the last Fully-Connected layer was reshaped into a 4×3 matrix, where each row contains the scores of a given character for each of the three labels. A softmax function was applied to the scores to represent the probability distribution of Yes, No, and DNK labels.
4. *CNN:* We additionally used a convolutional neural network (CNN) and the architecture of the CNN network is shown in (Extended Data Fig. 3a, Extended Data Table 8). We made the assumption that the information regarding each character’s appearance was encoded within a fixed time duration of the neural signal. Therefore, the windowed neural signal was treated as an image and fed into the CNN directly. The neural signal went through several convolution layers with LeakyReLU activations and Batch Normalization in between. The output of the last convolutional layer went through a Maxpooling layer and was subsequently flattened to a one-dimensional vector. Finally, the vector was fed into two sequential Fully-Connected layers with LeakyRelu activation and Batch Normalization in between. Similar to the LSTM network, the output of the last Fully-Connected layer was reshaped into a 4×3 matrix with rows and columns corresponding to the four characters and the three labels (Yes, No, and DNK) respectively. The scores were converted into probabilities after applying a softmax function. Here, too, the network architecture (e.g., number of layers and size) was optimized.
5. *Training, validation, and testing of the NN models*: The constructed dataset was randomly shuffled and split into training, validation, and testing sets. We used a 5-fold cross-validation method. The general procedure consisted of the following steps: a) We first split the randomly shuffled data into five groups (20% of the overall data in each group). b) For one fold of the cross validation, one group was picked as the test set, and from the remaining 80% of the data, we selected 87.5% as the training data (70% of the overall data) and 12.5% as the validation data (10% of the overall data). c) In each fold, the model was trained for up to 100 epochs (one epoch corresponds to one full iteration on the training data), and after each epoch, the model performance was evaluated on the validation set to infer its potential performance in the test set. This procedure is common practice to avoid over-fitting. d) Since each character’s label distribution was highly biased towards No, in order to ensure better predictions for each character, we selected the model with the best validation F_1_ score averaged within each character (note that accuracy is not a representative performance measure for imbalance datasets). e) For the selected model, the average performance on the test set in each fold was reported and summarized in Fig. 2, Extended Data Figure 2 and 3. It must be borne in mind that because each participant had a unique set of electrode locations, the models were trained and tested for each participant independently.
6. *Further details of the model*: The goal of training these networks in a supervised way was to predict the probability of a given character for each of the three labels (Yes, No, and DNK). Network weights were recursively updated using the Adam optimizer with backpropagation of the output’s computed loss. We initialized the weights with a random Gaussian distribution (mean=0, standard deviation=0.1) for the convolution layers, and a uniform distribution for Fully-Connected layers to better assist the gradient descent algorithm.
7. *Further quantifications of the model performance*: We used commonly used measures to quantify the performance of our models. It is worth noting that for measures (b-e), the reported results were averaged across the five folds (for each character, on the test set). After computing the number of true positives (TP), true negatives (TN), false positives (FP), and false negatives (FN), we also reported:

a. Normalized confusion matrix: TP and FN values were normalized by the total number of Yes labels, and TN and FP values were normalized by the total number of No labels.
b. Accuracy: (TP + TN)/(TP+TN+FP+FN) in Fig. 2, Extended Data Figure 3, Extended Data Table 7.
c. Recall: TP/(TP+FN) in Extended Data Table 7.
d. Precision: TP/(TP+FP) in Extended Data Table 7.
e. F-1 score: 2/(1/recall + 1/precision) in Fig. 2, Extended Data Figure 3, Extended Data Table 7, as well as in the text when identifying important brain regions in the decoding process. As mentioned previously, for unbalanced datasets, such as outs, F-score is a more realistic measure of the performance (as opposed to accuracy).
f. KLD loss: We computed the character-wise Kullback–Leibler divergence (KLD) loss to compare the difference between model predictions and ground truth. The mean of the four (character) losses was back-propagated to tweak the model’s weight. The loss associated with a DNK label was set to zero. Therefore, the back-propagated loss was only calculated on the average loss between Yes and NO entries in KLD loss for each character. Even though the label distribution was not even, no class weight was applied to the loss to balance the data. Furthermore, we also trained models with weighted loss over categories but got a similar performance. The fact that without designing handcrafted category weights and with unbalanced data, our models worked may also indicate that the vanilla KLD loss—together with complexity compatible architecture—was sufficient for mining essential patterns from a large enough dataset. The loss calculation is shown below:
KLD loss(y’, y) = Σ_c∈C_ y’(c) log(y’(c) / y(c)) where y’ are the model prediction, y are the ground truth, and c is the character label space.
g. *Comparisons of the NN model performances against a Naive Bayesian Model:* To further gauge our models’ performance, we compared our LSTM and CNN models against a Naive Bayesian model as quantified by accuracy, and F1-scores. For the Naive Bayesian model, we learned each character’s category distribution from training data, then made predictions over test data points following this distribution. One observation was that CNN and LSTM both achieved high performance with small variations in performance, which suggested that they all captured the essential information for decoding character footprints but may vary in the way of constructing the knowledge and making inferences. Additionally, we can conclude that without a structured and meaningful model for detecting and decoding, the F1-score would be far lower even with guesses following statistics.

### Determining the important brain regions in character decoding

To quantify the contributions of different regions to the decoding process, we performed multiple analyses:

1. *Region Knockout*: To evaluate a given brain region’s importance, we introduced a metric called knockout analysis. The method is analogous to occlusion sensitivity analysis, a tool commonly used for inspecting NN image classifiers^13^. Here, we perturbed the area of the input corresponding to each region (i.e., spiking activity of all of the neurons recorded from that region) by replacing it with zeros. Then, we re-evaluated the model performance on the perturbed data in the whole training dataset. The general procedure of this knockout analysis was performed by iterating over all regions (knocking out one region at a time) and computing accuracy, loss and F_1_ score for each character (within each participant). These performance metrics were averaged over models trained on each of the five folds. Furthermore, for the loss analysis, we subtracted the baseline loss (loss from the original model with intact inputs) from the loss obtained from the knockout perturbation for each region. Thus, a positive change in the loss indicates a lower decoding performance after a region has been knocked out. Lastly, to account for the different number of neurons recorded from each region, the change in the loss was normalized by the number of neurons (Fig.3a, b and Extended Data. Fig. 4). As previously mentioned, all of these analyses were done in a within-participant design manner.
2. *Electrode knockout*: To address whether the encoding correlations among the neurons within each region was contributing to the decoding performance, we explored the importance of each electrode following the same procedure described above (Fig. 3c and Extended Data Fig. 4) in the region knockout section and computed the same metrics (accuracy, F1-score, and loss). Concretely, we added up the change in the KLD loss following knocking out all the electrodes from a given region (one at a time, subtracting the baseline loss, but not normalizing by the number of neurons). Next, we subtracted this sum from the region knockout loss (after subtracting the baseline loss, but not normalizing by the number of neurons). A positive value would indicate that the information in a region, as a whole, is greater than simply the sum of the information from its constituents. The rationale behind a lack of normalization by the number of neurons in this particular step was that the number of neurons were the same on the two sides of the subtraction operation.
3. *Re-trained models:* To further verify the difference between important and non-important regions in the decoding process, within each participant, we employed an alternative approach. Here, we trained two separate models: 1) The first model was trained (and tested) using the neural activity from the regions that were deemed important from the region knockout analysis. 2) The second model was trained (and tested) using the neural activity from the remaining regions. The split was done such that the two models had comparable numbers of neurons as their inputs. Here, we computed the F1-scores for each model and compared the two across all characters and participants. The results, demonstrating that the model trained on important regions performed significantly better than the other model, are reported in the main text.

### Analyses concerning the memory test

As described in the Behavioral task section of the Methods, the memory test consisted of clip viewing and response time phases. For each participant, the neural data during the entire memory test was fed into the participant-specific model (that was trained and tested during movie viewing), and the model activation (probability of each character) was obtained as a function of time. Furthermore, we defined that a character is activated by the model in a given phase (i.e., clip viewing or response time) if model predictions went above 0.5 in that phase. The analyses presented in Fig.4 and Extended Data. Fig. 5 required several steps:

1. *Identifying the characters during each clip:* Here, character identification was done based on the clips such that, within each clip, we used the result of our semi-supervised character identification method, and defined a character as present if the number of their occurrences (positive detections) was above a threshold. We quantified the performance of the NN model predictions in decoding the characters during the clips while we allowed this threshold (percentage of time when the character was present in the clip) to vary. Similarly, we allowed the threshold for the size of the character in the frame (maximum=half of the frame) to vary and quantified the NN model performance respectively (Fig. 4b).
2. *Generalized Linear Models (GLM):* To examine what factors significantly contributed to the NN model activations during the memory test, we used a GLM method. Here, the character activations (averaged across the five folds and summed over a two-second period following the clip onset) were modeled as a function of 1) the presence or absence of the character in the clip; 2) whether the participant marked that clip as seen or not during response time; and 3) whether the clip belonged to the target category (same episode) or the lure category (another episode the participants had not seen). A similar analysis was done during a two-second period following the end of the clip as well as a two-second period prior to the participants’ responding time. The estimated coefficients and the corresponding p-values for each of these factors are reported in the main text.
3. *Character associations during the movie*: We defined a character occurrence association based on the scenes during the movie for which the timing information was provided by an independent study^7^. The occurrence association between two characters was defined as the sum of durations of scenes where both characters were present divided by the sum of durations of scenes where the conditioned character is present. To be more specific, the association between character c_i_ and c_j_ can be denoted as follows:

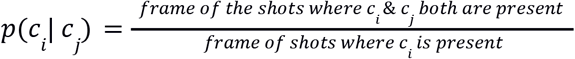

we will refer to this occurrence association (conditional occurrence probability), character associations.
4. *The conditional probability of model activation for characters:* We constructed the conditional activation probabilities of each pair of characters (P_model activation_(c_i_|c_j_)) over a given phase (clip viewing or response time) as the number of trials for which character i and character j are both activated divided by the number of trials where character j (the conditioned character) is activated by the model. Recall that, here, the activation is defined as the model predictions going above a threshold (0.5). Thus, a higher conditional probability indicates that the model predicts character i more frequently when character j is predicted. We generated two sets of 16 conditional activation probabilities between the four prominent characters for the clip viewing phase and the response time phase. These conditional activation probabilities were compared between clip viewing and response time phases using a signrank test, which revealed higher values during the response time (Fig. 4).
5. *The relationship between conditional activation probabilities and character associations during clip viewing:* Within each participant, we computed the (Spearman) correlation between the conditional probabilities of the model activation for the characters (computed during clip viewing; see item 4 above) and the character associations (computed over the movie duration; see item 3 above) after removing the diagonal elements from these matrices. The results are reported in Fig. 4 and the main text.

## Extended Data Figures and Tables

**Extended Data Figure 1.**
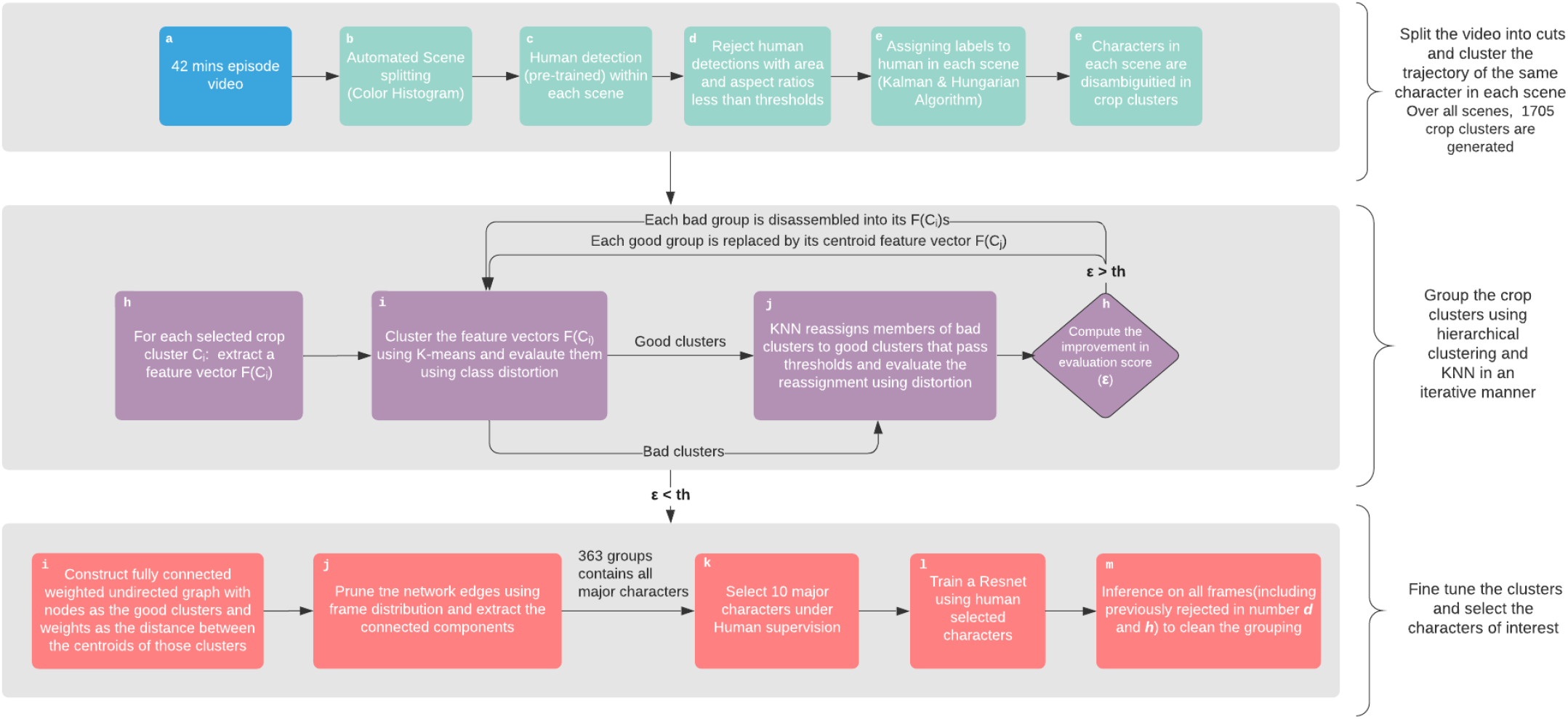
Flow chart of Semi-supervised Character Identification. We developed the semi-supervised framework to identify characters in the movie at the frame level. It consists of three main stages, requiring minimal human supervision but producing reliable ground truth labels to train our decoding models. In the first stage, we split the whole video into cuts based on the scenes (b), then created stable real-time identification of characters, i.e. crop clusters, based on human detection (c,d), and tracking and matching algorithm (e). In the second stage, we merged crop clusters purely based on facial features with k-means clustering (i) and KNN (j) iteratively.

**Extended Data Figure 2.**
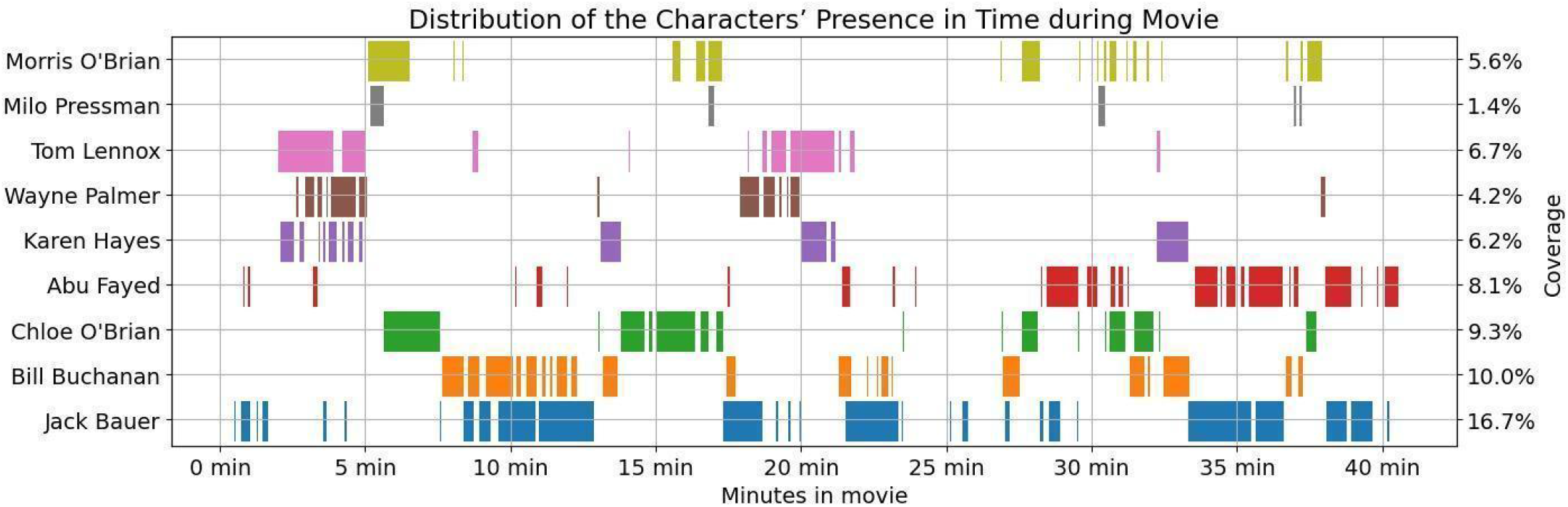
Distribution of the Characters’ Presence in Time during the Movie. The results from our semi-supervised character identification algorithm are shown. The presence of each of the nine characters (rows) in each frame of the movie (x-axis) is indicated by vertical lines. For our subsequent neural decoding analysis, we picked the four characters (bottom 4) that were most prominent and were present during different time points in the movie. For further details about the character distributions in the movie, see Extended Data Table 4. For visualization purposes, each vertical line has been dilated for 50 data points.

**Extended Data Figure 3.**
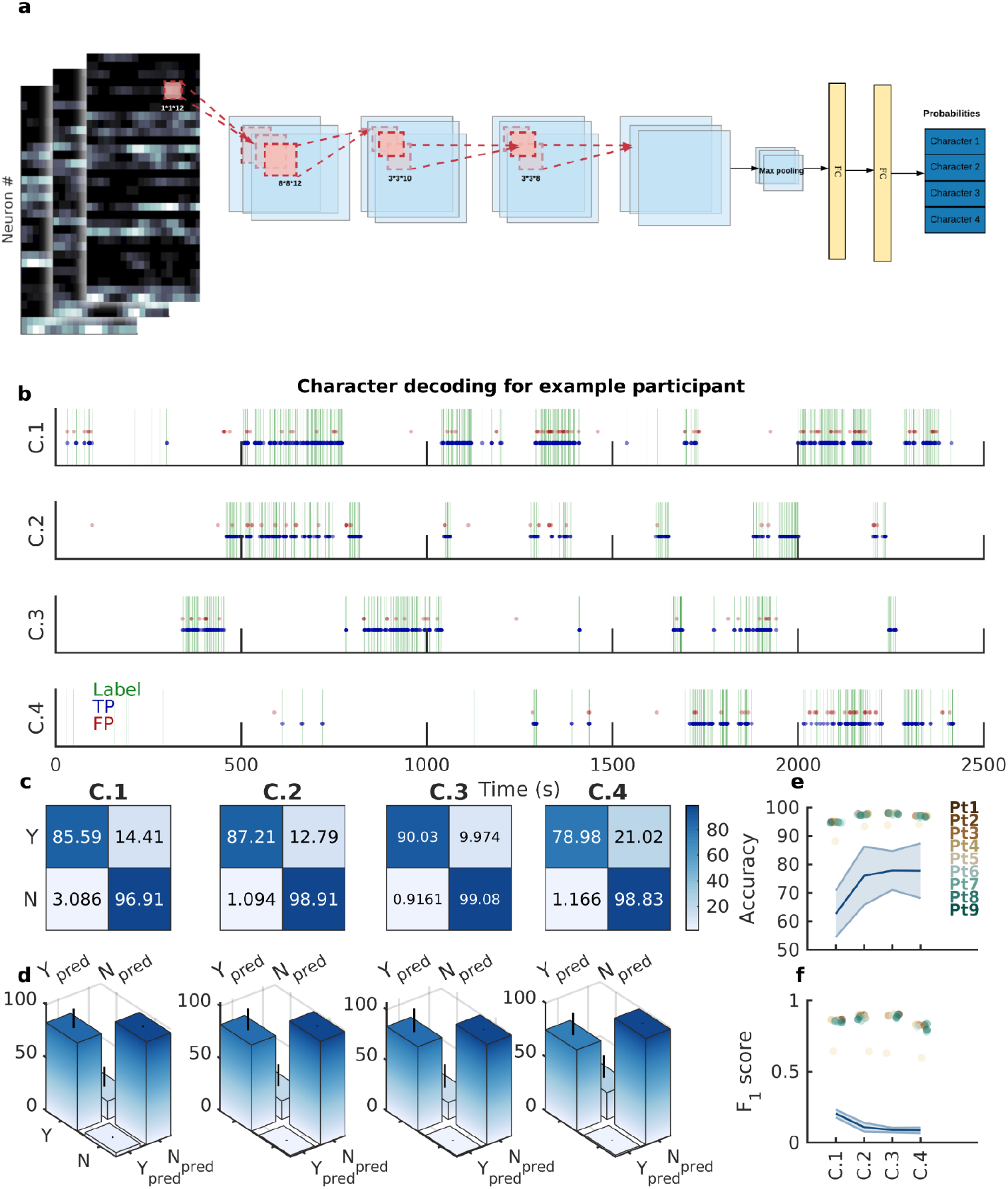
Convolutional Neural Network as an Alternative Architecture to Decode Characters from Neural Data (related to Fig. 2). **a.** The structure of the CNN network used for classification. Here, input data was the firing sequence of all neurons (colormap; x-axis: time, y-axis: neuron number; brighter shades correspond to higher firing) from a participant within a two-second window around each frame. Firing rate maps were passed through four CNN layers followed by two fully connected (FC) layers to output a probability distribution over the four main characters in that frame of the movie. **b.** Frame-by-Frame comparison of character labels generated by CV and CNN: The labels generated by the CNN and the computer vision algorithm for each movie frame are plotted as a function of time. The significant overlap between the two labels (a large number of true positives) illustrates the goodness of the decoding algorithm (A manual inspection of the time series output of the CNN model for each frame against the true character labels and noted that the model prediction shared high overlap with the true labels). **c.** In an example participant, the normalized confusion matrices for the binary classification task for all the four characters are shown. The large numbers on the diagonals (high TPR and TNR) of all the four matrices show that the CNN achieves high accuracy in decoding all the four characters. **d.** The distribution of the entries of the confusion matrix overall participants is shown as a bar plot (mean) with errorbar (std) for all four characters. The high mean and low standard deviation for the TPR and TNR values in all four matrices shows that the CNN achieves high accuracy in decoding all the four characters across participants. **e-f.** Accuracy (e) and F1-scores (f) for decoding each character are shown with each colored dot indicating different participants. The consistently high accuracy and F1-scores across participants indicate that the CNN generalizes well in this decoding task. The lines and shaded areas (mean±STD) indicate the performance of the chance model (obtained from shuffling labels) across all participants.

**Extended Data Figure 4.**
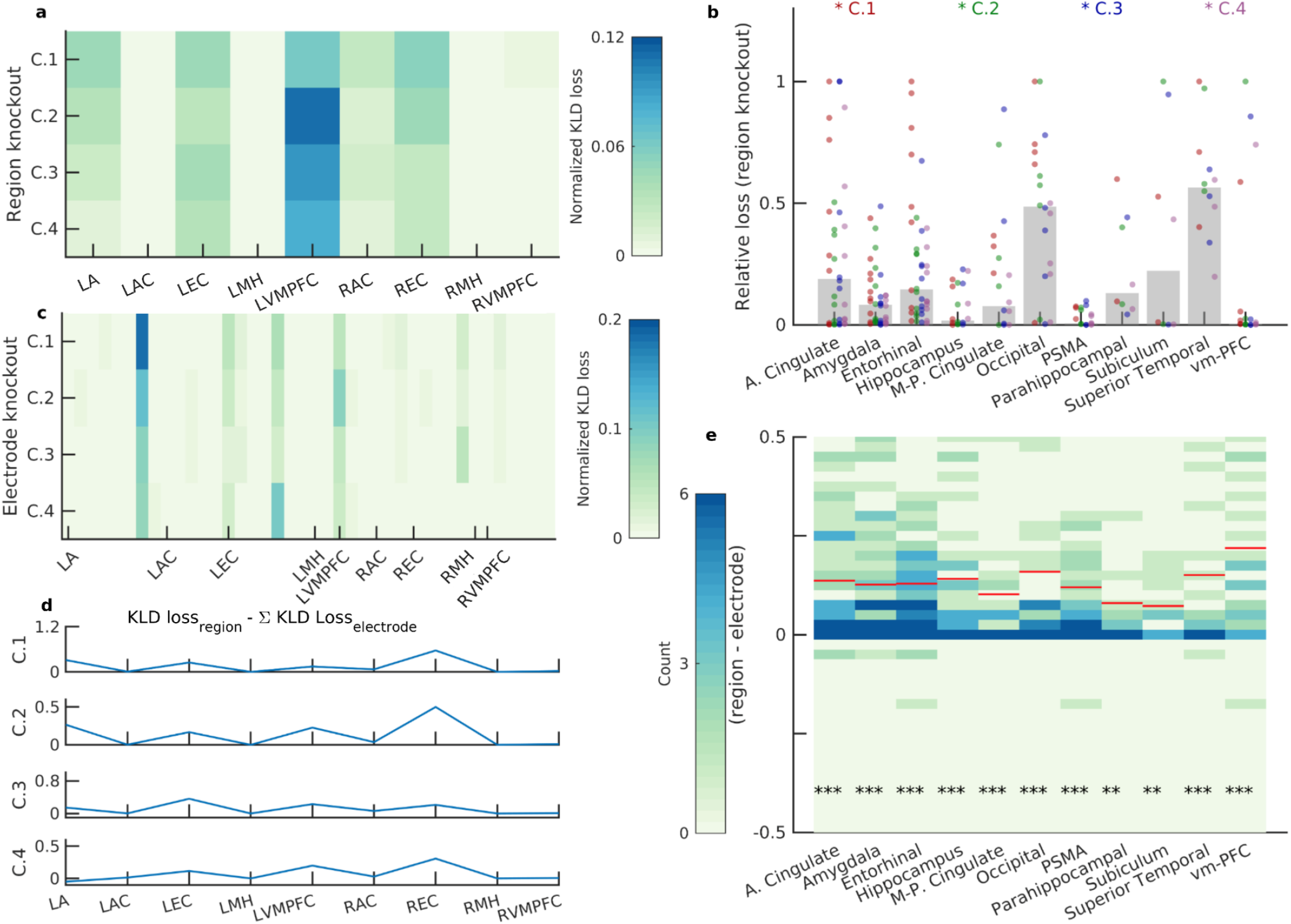
Identification of important regions in decoding characters using a CNN architecture (related to Fig. 3). **a.** The change of KLD loss for each character (row) after knocking out a given region (column) for one participant is shown (Region Knockout). The value is normalized by the number of neurons in that region and demonstrates how the model performance deteriorated when excluding the units recorded from that region. Important regions are those with higher KLD loss values. L and R correspond to the left and right hemisphere respectively. A: amygdala, AC: Anterior Cingulate, EC: Entorhinal Cortex, MH: Middle Hippocampus, VMPFC: Ventro-medial Prefrontal Cortex. **b.** The changes in KLD loss after knocking out regions are shown across participants. Different colored dots correspond to the change in KLD loss for different characters. Bars indicate the median value of the change in KLD loss after region knockout. The following regions resulted in the most notable losses in decoding performance: anterior cingulate (22.22, [10.12, 39.15]%), occipital (40.00, [19.12, 63.95]%), Subiculum (37.5, [8.52, 75.51]%), and superior temporal (66.67, [34.89, 90.08]%). Reported are the percentage of losses above 0.5 (as well as the binomial fit confidence intervals) **c.** The change in KLD loss for each character (row) after knocking out a given electrode (column) at a time is shown (Electrode Knockout) for an example participant (same as in a). Similar to the region knockout results in (a), the loss value is normalized by the numbers of units recorded on each electrode. **d.** The sum of the change in KLD loss following electrode knockout (all electrodes within a region) was subtracted from the change in KLD loss following region knockout. Shown are these values for the four different characters (rows) from an example participant (same as in a and c). Positive values indicate that knocking out a whole region deteriorates the model performance to a greater extent. **e.** When considering all regions from all participants, in most regions, the region knockout loss was greater than the sum of electrode knockout loss (each column, and its associated colormap, is the distribution of this measure and the red horizontal line indicates the median of the distribution for those that were significantly different from zero) as quantified by Wilcoxon signed-rank tests (*: p<0.05; **: p<0.01; ***: p<0.001).

**Extended Data Figure 5.**
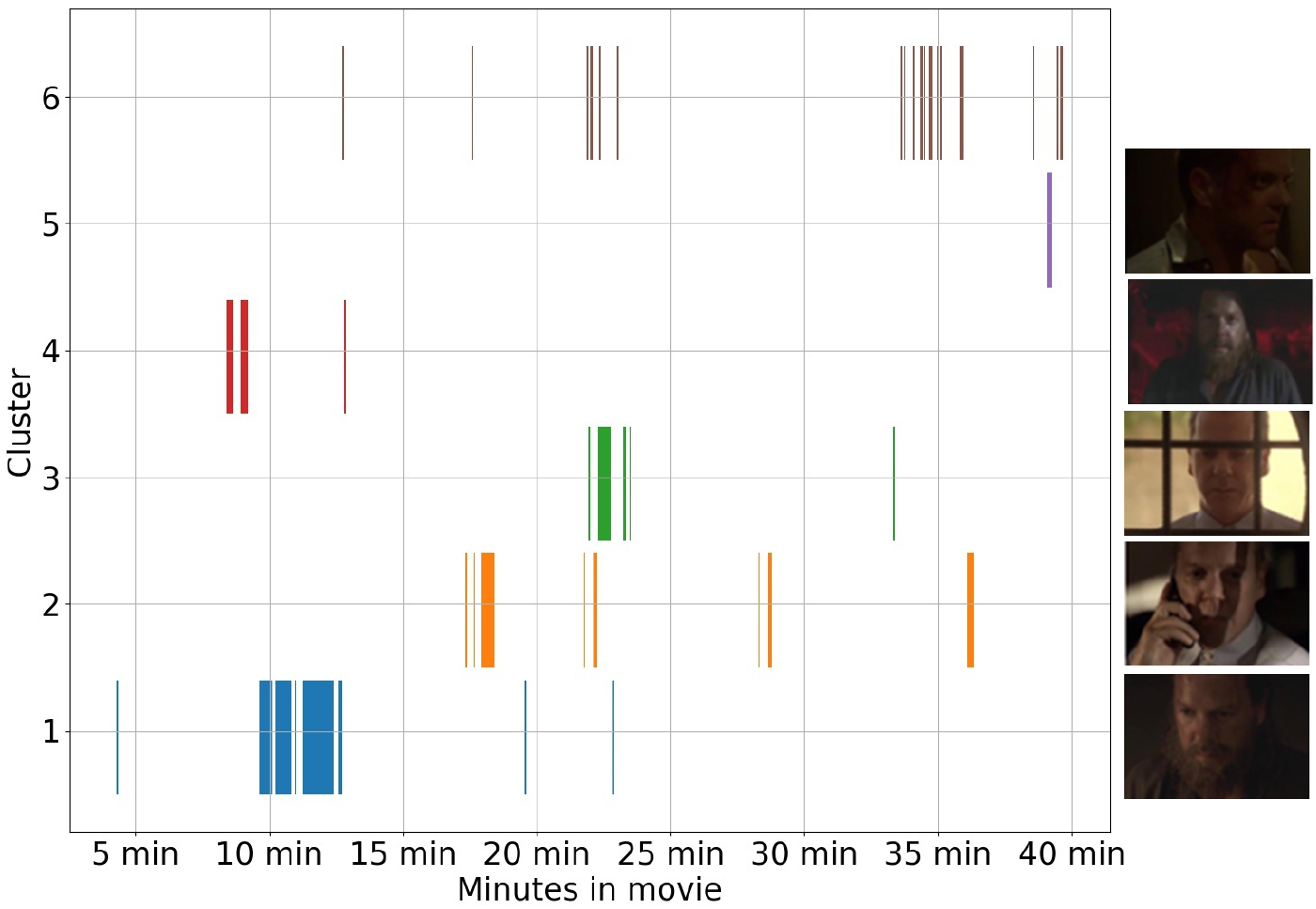
The Semi-supervised Character Identification Algorithm Generated Multiple Clusters for Each Character throughout the Movie. Our character identification method resulted in multiple clusters for a single character. Each row represents a cluster and the vertical lines indicate the frames in which a given cluster was identified. Note that each cluster corresponds to a visually different representation of the same character (snapshots on the right; cluster six refers to all other small clusters merging together).

**Extended Data Table 1.** Participant demographics. Demographics of the study participants (age, gender, and handedness) are presented.

| Demographics of Participants |  |  |  |
| --- | --- | --- | --- |
| Pt.<br>ID | Age | Handedness | Gender |
| 1 | 50 | R | F |
| 2 | 22 | L | F |
| 3 | 27 | R | F |
| 4 | 31 | R | M |
| 5 | 49 | R | F |
| 6 | 49 | R | M |
| 7 | 24 | R | F |
| 8 | 37 | L | M |
| 9 | 20 | R | M |

**Extended Data Table 2.**
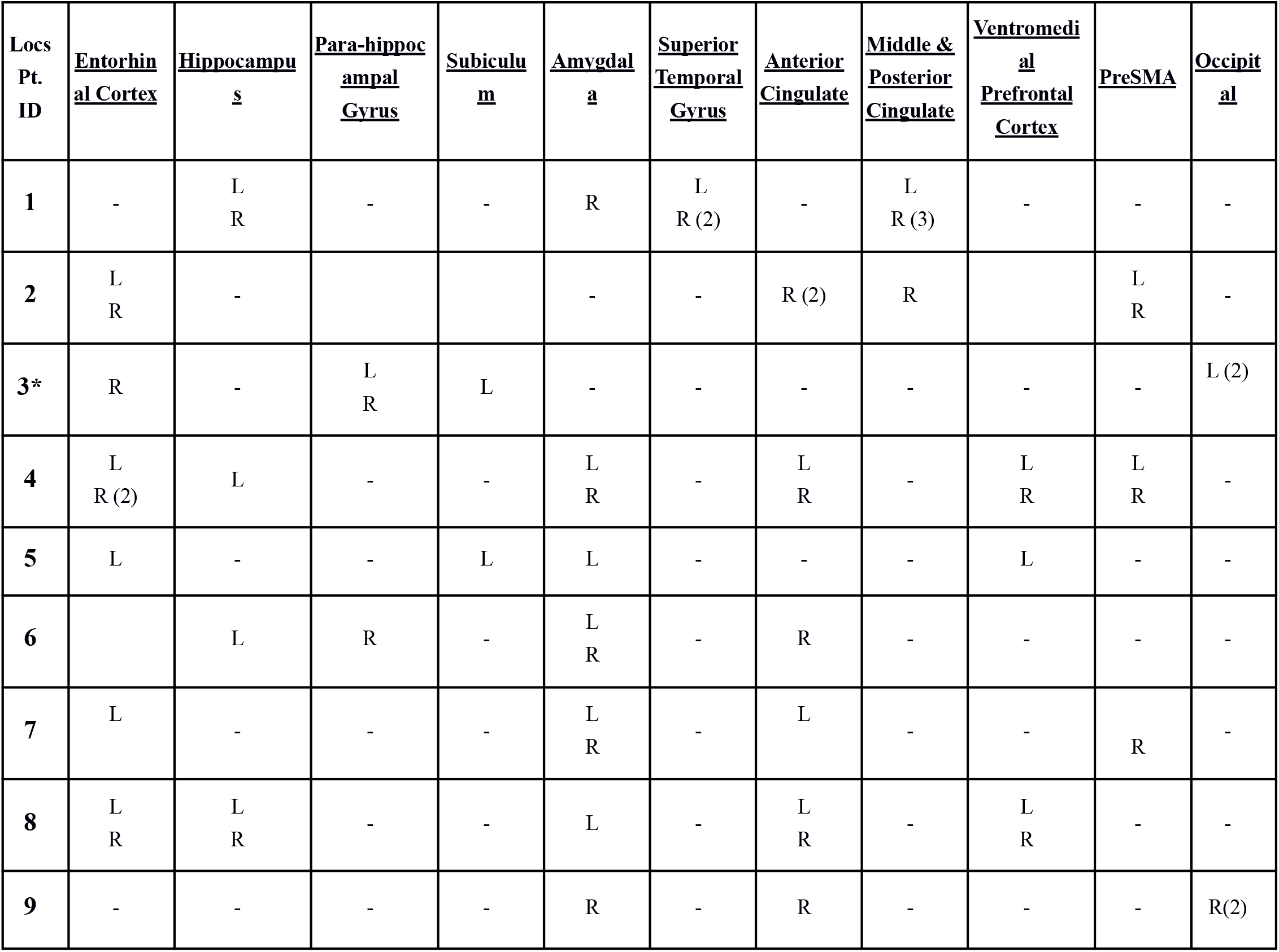
Electrode Localizations. Electrode locations are listed for each participant (rows). Columns indicate the electrode locations (categories that were used for group analysis) with R and L referring to the right and left hemispheres respectively. The number in parentheses indicates the number of electrodes within each hemisphere. *Participant 3 had units in the parietal region as well.

**Extended Data Table 3.** Number of Units Recorded in Each Participant.

| Participant ID | Number of Units |
| --- | --- |
| *1* | 56 |
| *2* | 53 |
| *3* | 49 |
| *4* | 84 |
| *5* | 8 |
| *6* | 18 |
| *7* | 29 |
| *8* | 60 |
| *9* | 28 |

**Extended Data Table 4.** Distribution of the Character Labels during the Movie Generated by the Semi-supervised Framework. This table shows the percentage of each label for the four main characters detected by our semi-supervised character identification framework on a frame level in the movie. Label Yes (No) was defined as when the character was (not) present at the exact time step (frame). Label DNK (Do Not Know) was introduced as a mechanism to particularize the training set (Methods). The characters were present in less than 20% of the frames across the movie and, thus, it must be borne in mind that the training data used for further neural decoding model was heavily skewed and appropriate measures were taken to remedy this issue. The percentage of DNK was around 2% of the total training samples, which indicates that the dropping of the data sample would not affect the total information provided to the model.

|  | Yes | No | DNK |
| --- | --- | --- | --- |
| C.1 | 17.96 | 80.25 | 1.826 |
| C.2 | 10.73 | 85.73 | 3.531 |
| C.3 | 10.01 | 89.76 | 2.230 |
| C.4 | 8.683 | 89.59 | 1.720 |

**Extended Data Table 5.**
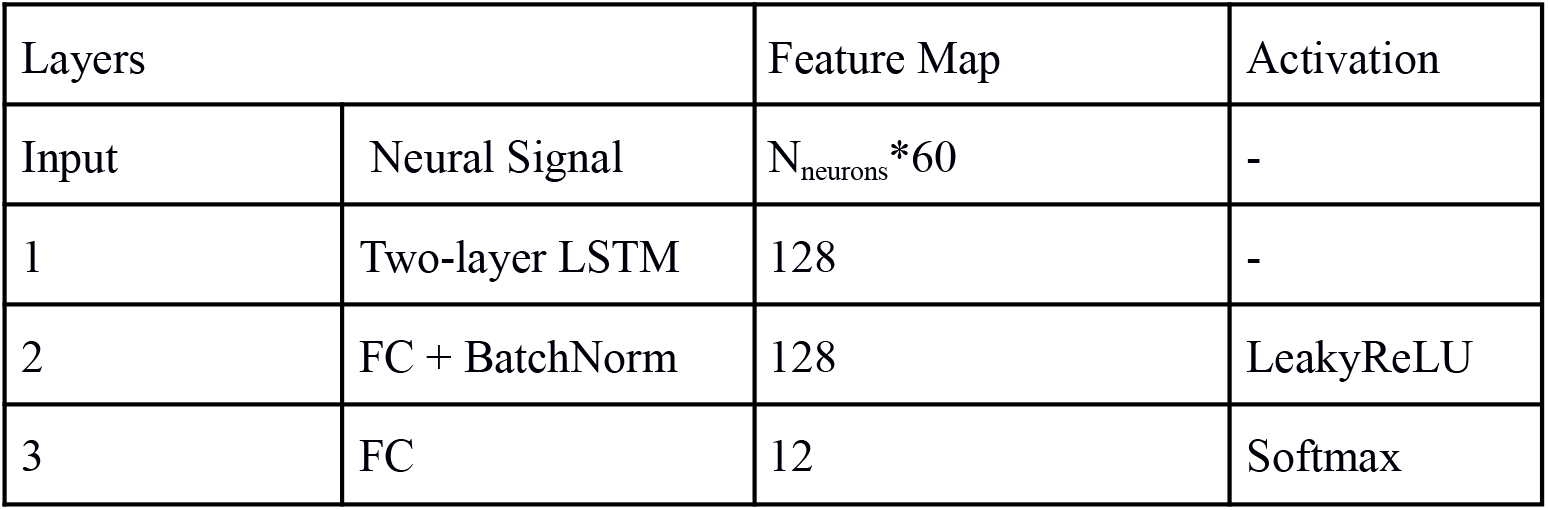
LSTM Architecture. The input to the LSTM model consisted of the firing rate of all neurons from a participant in time (2 seconds around each frame with 60 time-steps). This was first fed into a two-layer LSTM followed by a fully connected layer (FC) and a Batch Normalization layer (BatchNorm) with LeakyReLU as the activation function. This was further processed by a connected layer followed by a softmax operation to output the confidence scores for three labels (Yes, No, DNK) for each character.

**Extended Data Table 6.** Comparing NN Model Performances against Baseline (Naive Bayes). For each participant (columns), average performance (as quantified by F1-scores) is reported for three different methods used to decode the visual presence of characters using the neural data. Our main methods, the LSTM and the complimentary CNN architectures, fared much (on average 7 times) better when compared against the performance of a Naive Bayes method, which was used as a baseline model.

|  | Pt. 1 | Pt. 2 | Pt. 3 | Pt. 4 | Pt. 5 | Pt. 6 | Pt. 7 | Pt. 8 | Pt. 9 |
| --- | --- | --- | --- | --- | --- | --- | --- | --- | --- |
| **NB** | 0.1160 | 0.1144 | 0.1158 | 0.1154 | 0.1180 | 0.1158 | 0.1148 | 0.1137 | 0.1151 |
| **LSTM** | 0.8784 | 0.8766 | 0.8788 | 0.8772 | 0.7597 | 0.8536 | 0.8692 | 0.8793 | 0.8647 |
| **CNN** | 0.8738 | 0.8737 | 0.8633 | 0.8755 | 0.6272 | 0.8620 | 0.8400 | 0.8702 | 0.8483 |

**Extended Data Table 7.**
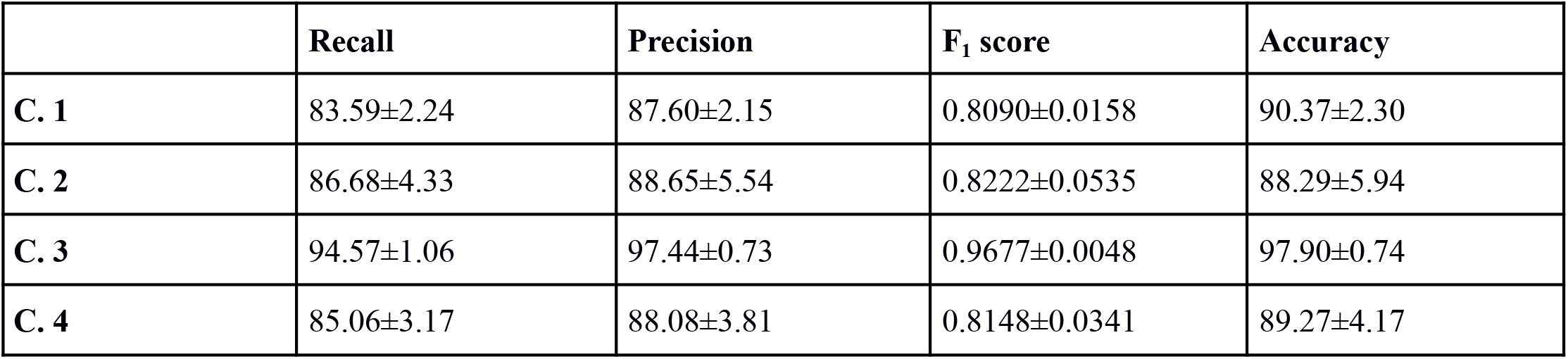
Further Quantifications of the LSTM Model Performance in Decoding Characters. In addition to the LSTM model F1-scores and accuracy values shown for all participants in Fig. 2e and Fig. 2f, here we report mean±STD values of other measures such as recall and precision to quantify model performance in decoding each of the characters (rows).

|  | Recall | Precision | F<sub>1</sub> score | Accuracy |
| --- | --- | --- | --- | --- |
| C. 1 | 83.59±2.24 | 87.60±2.15 | 0.8090±0.0158 | 90.37±2.30 |
| C. 2 | 86.68±4.33 | 88.65±5.54 | 0.8222±0.0535 | 88.29±5.94 |
| C. 3 | 94.57±1.06 | 97.44±0.73 | 0.9677±0.0048 | 97.90±0.74 |
| C. 4 | 85.06±3.17 | 88.08±3.81 | 0.8148±0.0341 | 89.27±4.17 |

**Extended Data Table 8.** CNN Architecture. The CNN model was used as an independent confirmation of the LSTM model results. Here, too, the windowed neural signal (firing rate of all neurons from a participant during a two-second interval around each frame) was the input that underwent several convolution layers with LeakyReLU activations and Batch Normalization (BatchNorm) layers in between. The output of the last convolutional layer went through a MaxPooling layer and was then flattened to a one-dimensional vector. This one-dimensional vector was concatenated with the flattened region tag embedding corresponding to each region in order to incorporate more information into the model. Finally, the concatenated vector was fed into two sequential Fully Connected (FC) layers with LeakyReLU activation and a Batch Normalization in between. The output of the last Fully Connected layer was reshaped into a 4*3 matrix corresponding to the confidence scores of the three labels (Yes, No, DNK) for each of the four main characters.

| Layers |  | Feature Map | Kernel Size | Stride | Activation |
| --- | --- | --- | --- | --- | --- |
| Input | Neural Signal | $N_{\text{neurons}} * 60 * 1$ | - | - | - |
| 1 | Conv + BatchNorm | $n_1 * 60 * 12$ | 1*1 | 1 | LeakyReLU |
| 2 | Conv + BatchNorm | $n_2 * 12 * 12$ | 8*8 | 1 | LeakyReLU |
| 3 | Conv + BatchNorm | $n_3 * 10 * 10$ | 3*3 | 1 | LeakyReLU |
| 4 | Conv + BatchNorm | 8 | 3*3 | 1 | LeakyReLU |
| 5 | MaxPooling | - | 3*3 | 3 |  |
| 6 | Flatten | Variable (see below)* | - | - | - |
| 7 | FC + BatchNorm | 64 | - | - | LeakyReLU |
| 8 | FC | 12 | - | - | Softmax |

## Footnotes

* Given that each participant had a different number of contacts, the number is variable from participant to participant.

## References

1. Abbas, Q., Ibrahim, M. E. A. & Jaffar, M. A. A comprehensive review of recent advances on deep vision systems. Artif. Intell. Rev. 52, 39–76 (2019).

2. Ranjan, R. et al. Deep Learning for Understanding Faces: Machines May Be Just as Good, or Better, than Humans. IEEE Signal Process. Mag. 35, (2018).

3. Quiroga, R. Q., Reddy, L., Kreiman, G., Koch, C. & Fried, I. Invariant visual representation by single neurons in the human brain. Nature 435, 1102–1107 (2005).

4. Quiroga, R. Q., Reddy, L., Koch, C. & Fried, I. Decoding visual inputs from multiple neurons in the human temporal lobe. J. Neurophysiol. 98, 1997–2007 (2007).

5. Gelbard-Sagiv, H., Mukamel, R., Harel, M., Malach, R. & Fried, I. Internally generated reactivation of single neurons in human hippocampus during free recall. Science (80-.). 322, 96–101 (2008).

6. Quiroga, R. Q. Concept cells: the building blocks of declarative memory functions. Nat. Rev. Neurosci. 13, 587–597 (2012).

7. Tang, H. et al. Predicting episodic memory formation for movie events. Sci. Rep. 6, 30175 (2016).

8. Quiroga, R. Q., Nadasdy, Z. & Ben-Shaul, Y. Unsupervised Spike Detection and Sorting with Wavelets and Superparamagnetic Clustering. Neural Comput. 16, 1661–1687 (2004).

9. Ison, M. J. J., Quian Quiroga, R., Fried, I., Quian Quiroga, R. & Fried, I. Rapid Encoding of New Memories by Individual Neurons in the Human Brain. Neuron 87, 220–230 (2015).

10. Suthana, N. A. et al. Specific responses of human hippocampal neurons are associated with better memory. Proc. Natl. Acad. Sci. U. S. A. 112, 10503–8 (2015).

11. Redmon, J. & Farhadi, A. YOLOv3: An Incremental Improvement. (2018). Available at: http://arxiv.org/abs/1804.02767. (Accessed: 12th August 2020)

12. Hochreiter, S. & Schmidhuber, J. Long Short-Term Memory. Neural Comput. 9, 1735–1780 (1997).

13. Zeiler, M. D. & Fergus, R. Visualizing and Understanding Convolutional Networks. in European Conference on Computer Vision (ECCV) 818–833 (Springer, Cham, 2014).

14. Davachi, L. Item, context and relational episodic encoding in humans. Current Opinion in Neurobiology 16, (2006).

15. Eichenbaum, H., Yonelinas, A. P. & Ranganath, C. The medial temporal lobe and recognition memory. Annual Review of Neuroscience 30, (2007).

16. Squire, L. R., Stark, C. E. L. & Clark, R. E. The medial temporal lobe. Annual Review of Neuroscience 27, 279–306 (2004).

17. Bettencourt, L. M. A. The Rules of Information Aggregation and Emergence of Collective Intelligent Behavior. Top. Cogn. Sci. 1, (2009).

18. Favila, S. E., Lee, H. & Kuhl, B. A. Transforming the Concept of Memory Reactivation. Trends Neurosci. (2020). doi:10.1016/J.TINS.2020.09.006

19. Manning, J. R., Sperling, M. R., Sharan, A., Rosenberg, E. A. & Kahana, M. J. Spontaneously reactivated patterns in frontal and temporal lobe predict semantic clustering during memory search. J. Neurosci. 32, 8871–8 (2012).

20. Miller, J. F. et al. Neural activity in human hippocampal formation reveals the spatial context of retrieved memories. Science 342, 1111–4 (2013).

21. St-Laurent, M., Abdi, H. & Buchsbaum, B. R. Distributed Patterns of Reactivation Predict Vividness of Recollection. J. Cogn. Neurosci. 27, 2000–2018 (2015).

22. Gordon, A. M., Rissman, J., Kiani, R. & Wagner, A. D. Cortical Reinstatement Mediates the Relationship Between Content-Specific Encoding Activity and Subsequent Recollection Decisions. Cereb. Cortex 24, 3350–3364 (2014).

23. Kahana, M. J., Howard, M. W. & Polyn, S. M. Associative retrieval processes in episodic memory. in Learning and Memory: A Comprehensive Reference (2007). doi:10.1016/B978-012370509-9.00185-6

24. M Aghajan., Z. et al. Theta Oscillations in the Human Medial Temporal Lobe during Real-World Ambulatory Movement. Curr. Biol. 27, 3743–3751.e3 (2017).

25. Wojke, N., Bewley, A. & Paulus, D. Simple online and realtime tracking with a deep association metric. in 2017 IEEE International Conference on Image Processing (ICIP) 3645–3649 (IEEE, 2017). doi:10.1109/ICIP.2017.8296962

26. Schroff, F., Kalenichenko, D. & Philbin, J. FaceNet: A unified embedding for face recognition and clustering. in Proceedings of the IEEE Computer Society Conference on Computer Vision and Pattern Recognition 815–823 (2015). doi:10.1109/CVPR.2015.7298682

27. He, K., Zhang, X., Ren, S. & Sun, J. Deep Residual Learning for Image Recognition. arXiv:1512.03385 [cs.CV] (2016).

28. Tampuu, A., Matiisen, T., Ólafsdóttir, H. F., Barry, C. & Vicente, R. Efficient neural decoding of self-location with a deep recurrent network. PLoS Comput. Biol. 15, (2019).

29. Castellano, B. PySceneDetect v0.5.5 Manual — PySceneDetect v0.5.5 documentation. Available at: https://pyscenedetect-manual.readthedocs.io/en/latest/. (Accessed: 27th April 2021)

